# Endometriosis lesions are oligoclonal structures derived from the normal endometrium

**DOI:** 10.64898/2026.02.25.708037

**Authors:** Sigurgeir Ólafsson, Ásgeir Ö. Arnþórsson, Kirsten Kübler, Ásgeir Sigurðsson, Helga Sigrún Gunnarsdóttir, Hákon Jónsson, Bergrún Ásbjörnsdóttir, Louise le Roux, Jóna Sæmundsdóttir, Valgerdur Steinthorsdottir, Guðmundur Norddahl, Ingileif Jónsdóttir, Jón Gunnlaugur Jónasson, Droplaug Magnúsdóttir, Ólafur Magnússon, Anna M. Jónsdóttir, Ragnheiður O. Árnadóttir, Kári Stefánsson

**Author notes:** Please address correspondence to.

## Abstract

Endometriosis is characterized by the presence of endometrium-like tissue outside the uterus. The origin of this ectopic tissue is debated, with leading theories including retrograde menstruation and embryonic remnants. Using somatic mutations as markers, we show that endometriosis lesions consist of unrelated clones of epithelial cells, with stromal cells being distinct, and that lesions at different body sites have different sets of clones. We observed that mutation burdens, signatures and driver landscapes are similar in cells from lesions and normal endometrium and the distribution of somatic mutations along the length of chromosomes is consistent with uterine origin. Furthermore a large-scale screen of the normal endometrium of endometriosis patients identified clones that are ancestral to endometriosis, indicating that the ectopic tissue in endometriosis originates from the normal endometrium.

## Main text

Endometriosis is a chronic condition characterized by the presence of endometrial tissue, epithelium and stroma, outside the uterus. It is associated with pelvic pain and infertility and is thought to affect as many as 10% of women of reproductive age (*1, 2*). Lesions present as any of three anatomical subtypes: (i) Superficial endometriosis, where lesions are confined to the pelvic area; (ii) deep infiltrating endometriosis, where lesions are found deep within pelvic structures and/or invading visceral organs; or (iii) as endometriomas, which are large cysts on the ovaries filled with dark brown endometrial fluid and lined on the inside with a thin sheet of endometrial cells. Patients often present with multiple lesions distributed throughout the pelvis and beyond (*1, 2*).

The origins of the ectopic tissue continue to be debated (*3, 4*). In the 1920s, John A. Sampson put forth his theory of retrograde menstruation, which maintains that endometrial cells, epithelium and stroma, refluxes through the fallopian tubes into the peritoneal cavity to implant on the pelvic surfaces (*5*). While intellectually compelling, this theory is supported by limited functional evidence in humans, and several competing theories have been proposed. These include the theory of coelomic metaplasia (*6, 7*), which states that endometrial cells arise through transformation of mesothelial cells in the peritoneum, and the Mullerian remnants theory, which states that lesions arise from cells which failed to properly differentiate or were displaced during fetal development (*8, 9*). Finally, the stem cell recruitment theory states that endometriosis lesions originate from stem cells travelling through the angiolymphatic circulation (*10*), but opinions are divided as to whether these stem cells would be of uterine origin or derived from other stem-cell niches, in particular the bone marrow (*11, 12*).

In addition to the unclear etiology of the lesions, the mechanism of endometriotic lesion dissemination within the body is incompletely understood (*4*). Specifically, do lesions spread through shedding of cells from an initial founder lesion, reminiscent of cancer metastases, or are lesions independently seeded by distinct progenitor cells?

All cells of the body continuously accrue somatic mutations throughout life (*13, 14*). The glands of the normal endometrium, which are structures of clonal cells, accumulate ∼30 mutations per cell per year (*15*). These mutations are naturally occurring lineage tracing markers, with mutations shared by different cells implying a common progenitor. While the majority of somatic mutations are thought to be functionally neutral, cancer driver mutations are also detected within normal tissues.

The normal endometrium in particular, is rich in mutations in genes commonly mutated in endometrial adenocarcinomas, including *PIK3CA, KRAS, ARHGAP35* and others (*15*–*17*). Driver mutations have also been reported at a high prevalence in endometriotic lesions (*17*–*19*), although their role in the etiology of the disease remains unclear.

Here we used laser capture microdissection to isolate individual endometrial glands from endometriotic lesions and patient-matched normal endometrium (*15, 20*). We performed whole-genome sequencing (WGS) of individual endometrial glands to identify the somatic mutations present and build phylogenetic trees to reveal the developmental histories and relationships between lesions and to compare the somatic mutation landscape between normal tissue and endometriotic lesions.

## Results

### Somatic mutations in endometriosis mirror those found in normal endometrium

We obtained tissue biopsies of endometriotic lesions and matched normal endometrium from 25 women undergoing laparoscopic surgery as part of their treatment for endometriosis at Landspitali University Hospital in Iceland between 2022 and 2024 (Methods, Table S1). We used laser capture microdissection to isolate samples of a few hundred endometrial or stromal cells for WGS (*15, 20*) (Figure 1A, Methods). The dataset comprises 403 microdissections in total; 315 microdissections of epithelium from endometriosis biopsies, 3 samples of adjacent stroma, 3 from adenomyosis and 85 glands from matched normal endometrium sequenced to a median depth of 19X (range 5-106X, Figure S1, Table S2). After filtering and quality control (Methods), 213,280 substitutions and 11,071 indels were used in the analyses described below.

**Figure 1.**
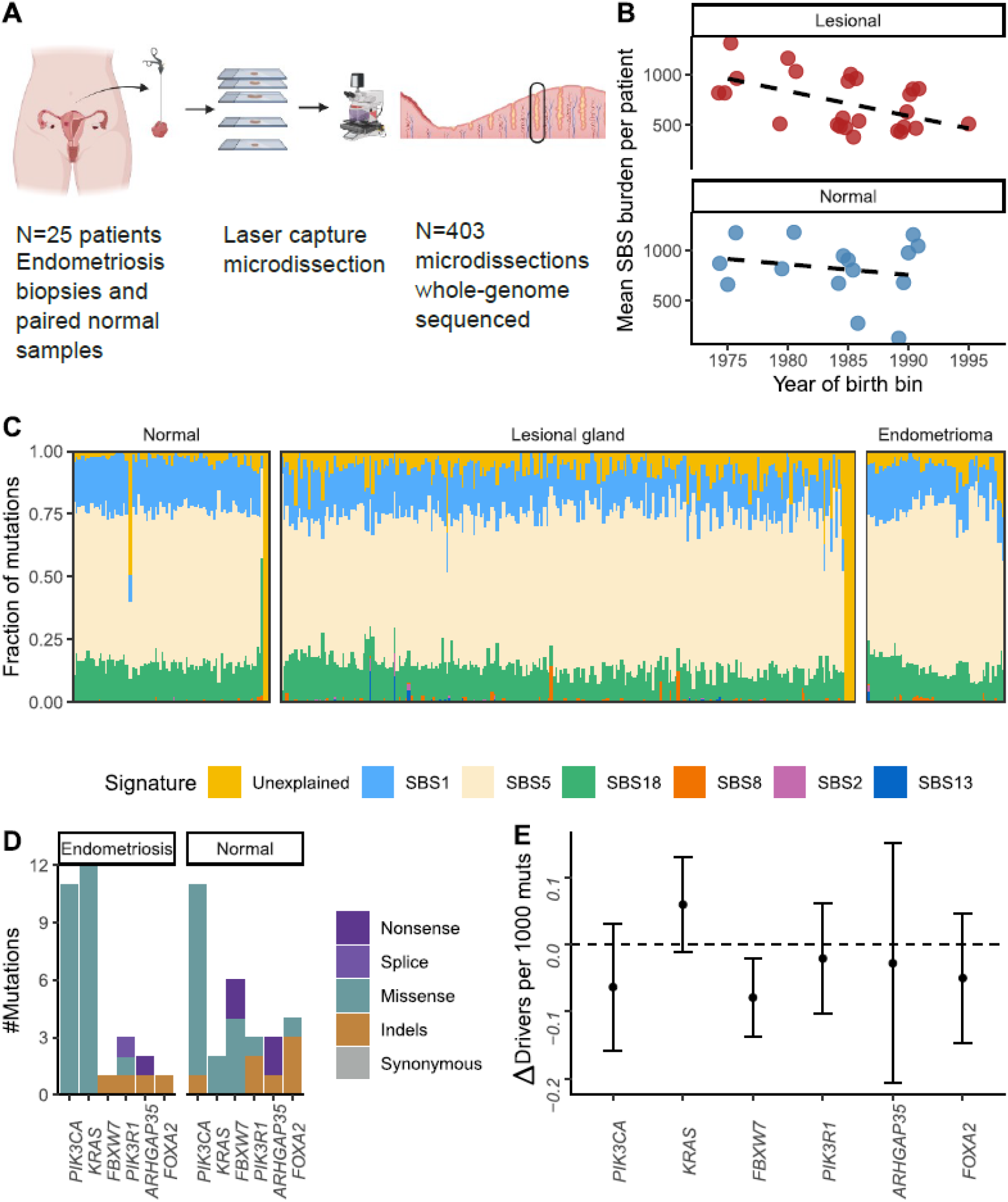
Somatic mutations in endometriosis and normal endometrium. A) An overview of the cohort and sampling. B) Mutation burden in endometrial glands isolated from lesional and normal tissue in 5-year year-of-birth intervals. C) Substitution mutational signatures identified in microdissected samples. Each bar represents one microdissected sample. D) An overview of the mutations identified in each of the genes showing evidence of positive selection in our cohort. E) The change in the number of driver mutations observed per 1000 mutations between lesional and normal endometrial cells. Error bars represent 95% confidence intervals.

We fit a linear mixed-effects model to jointly estimate the effects of the age of the donor and lesional status after correcting for sequencing coverage, median variant allele fraction (VAF) and non-random sampling (*21*) (Methods). We estimate a yearly increase of 23.3 mutations per cell (7.1-39.5 95% CI, P=0.0078, Likelihood ratio test (LRT), Figure 1B) and found that lesional samples do not have a mutation burden different from normal (P=0.62, LRT). We used a Bayesian hierarchical Dirichlet process to extract COSMIC mutational signatures for all samples. In line with previous work (*13, 15*), we found the mutation spectra of all samples to be dominated by SBS1, SBS5 and SBS18. These signatures contributed an equal number of mutations in cells from lesional and normal tissues, suggesting that epithelial cells in endometriotic lesions are subject to the same mutational processes as the cells of the normal endometrium and to a similar magnitude (Figure 1C).

We used the dNdScv software (*22*) (v. 0.0.1.0) to identify genes where the ratio of the mutation rate at non-synonymous sites (dN) and synonymous sites (dS) is greater than 1, indicating positive selection of non-synonymous mutations. Combining mutations from lesional and normal biopsies, six genes showed evidence of positive selection after Benjamini-Hochberg correction for multiple testing (Figure 1D, Table S3). No additional genes could be identified in analyses stratified by lesional status (Table S3). Taking into account differences in the number of samples and clonal structure between lesional and normal biopsies (Methods), no gene was differentially mutated in lesional samples compared with normal endometrium after correction for multiple testing. *KRAS* mutations have been previously implicated in endometriosis (*17*–*19*) and are specially highlighted in Figures S4-S27.

*KRAS* hotspot mutations were found in 50/315 (16%) of microdissections from endometriosis compared with 3/85 (4%) normal glands. However, the *KRAS* mutations in endometriosis often preceded clonal expansions and so only represent 12 independent mutation events. Thus, although we observed 4-times as many hotspot mutations in *KRAS* in lesional compared with normal samples, this difference is not statistically significant (Figure 1E, P=0.1, likelihood ratio test of missense mutations, Coselens (*23*)) after accounting for differences in clonal structure between samples.

Taken together, these results show that somatic mutations in epithelial cells in endometriosis mirror those found in normal endometrium.

### Individual lesions are derived from multiple distinct cells

To gain insights into the cellular histories of endometriotic lesions, we reconstructed phylogenetic trees (MPBoot software (*24*) (v. 1.1.0)) for cells isolated from each patient (Figure 2, Figures S4-S27). Inspecting the trees from all 25 patients, three broad insights emerge from our data:

**Figure 2.**
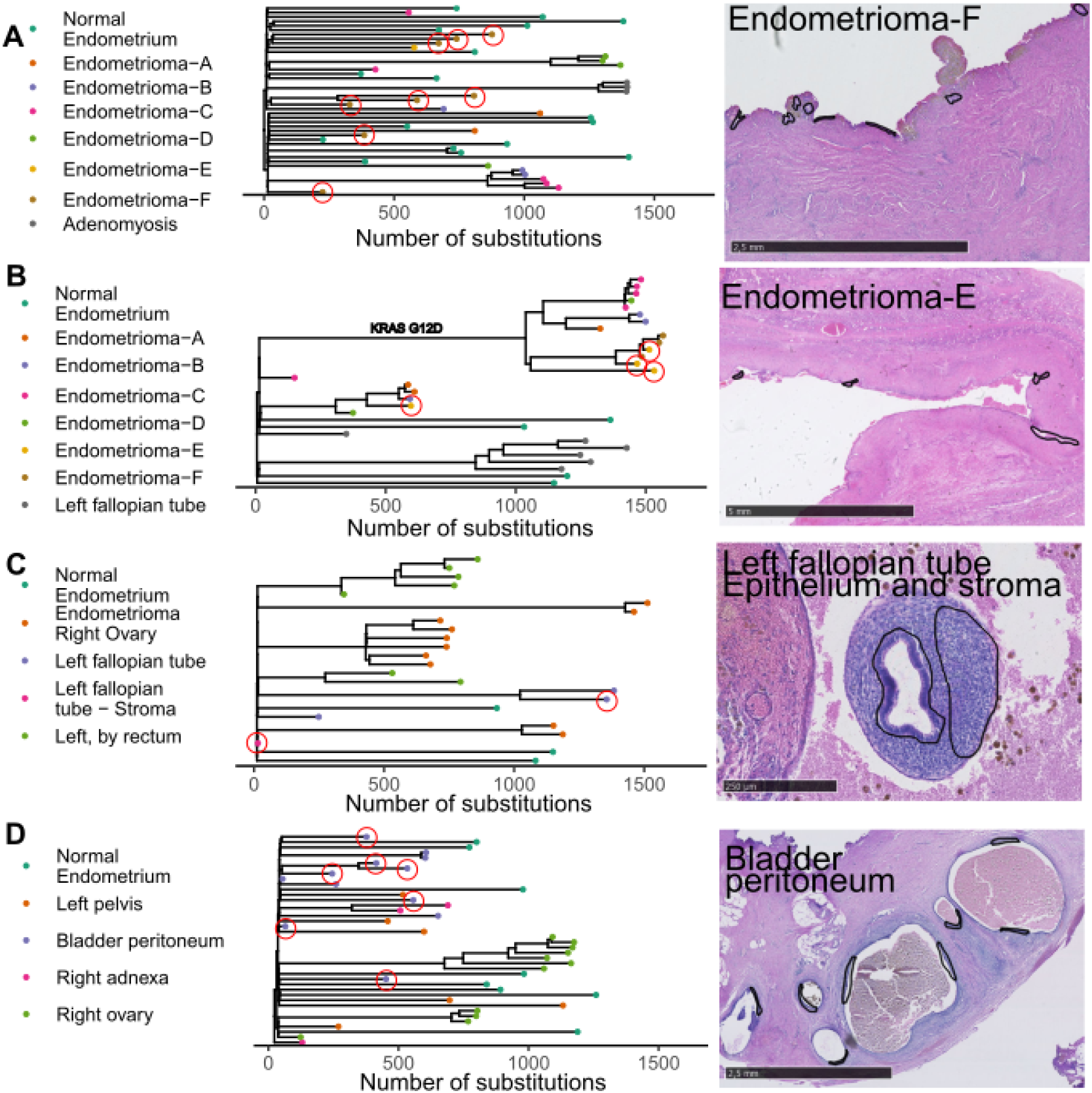
Examples of phylogenetic trees from endometriosis patients. Tips of trees circled in red are microdissections marked with circles in black in the H&E tissue images to the right. The x-axis is the same across all panels. A) Patient 3, presenting with endometrioma and adenomyosis. Multiple independent cell clones are highlighted within a 3-4mm wide section of the endometrioma. B) Patient 30, presenting with endometrioma, which was sectioned into slices separated by 3-4 mm of tissue, creating biopsies A-F. A large clone spanning all biopsies and carrying a *KRAS* G12D mutation was identified but other independent clones are also found within the endometrioma, showing oligoclonal origin. C) Patient 37, presenting with endometrioma as well as superficial and deep-infiltrating lesions. Cells isolated from different lesions do not share mutations. The histopathological image shows an example of a gland and adjacent stroma. No mutations could be detected in the stroma sample, most likely due to polyclonal composition. D) Patient 18, presenting with superficial endometriosis lesions on multiple body sites. No sharing of mutations is observed between sites of the body. The highlighted lesion manifests as rows of cyst-like structures on the peritoneum of the bladder. Each cyst is derived from an independent cell, sharing no mutations with its neighbor.

First, individual endometriosis lesions typically comprise multiple unrelated clones of epithelial cells. This shows that epithelial cells within a lesion are generally not seeded by a single cell but rather by multiple cells which have diverged from one another during embryonic development and thus share few mutations in common. An example is shown in Figure 2A. Multiple microdissections from the same 2-3 mm strip of endometrioma (Figure 2A-right) were sequenced and found to share few-to-no mutations in common. These cells are also unrelated to cells isolated from other biopsies of the same endometrioma, revealing the endometrioma to be a polyclonal structure.

Figure 2B shows a second example of this. An endometrioma was sectioned, with samples taken every 3-4 mm, yielding biopsies A through F. While in this case a large clone (carrying a *KRAS* G12D mutation) was found to span all the biopsies, other independent clones were also found within the endometrioma. The endometrioma clones are also unrelated to a superficial lesion on the fallopian tube (gray) displaying a dichotomous tree structure. This superficial lesion exemplifies how samples from the same clade of the tree are sometimes separated by dozens to hundreds of mutations. The coalescent events defining these clades are rarely polytomies but rather display a bifurcation pattern consistent with continued local growth of the clone. This branching structure suggests that epithelial cells continue to expand locally within the host tissue post seeding.

The second insight emerging, as suggested by previous work (*25*), is that stroma cells from endometriosis lesions are unrelated to the epithelial cells. We sequenced 3 samples of stroma directly adjacent to glands of epithelial cells of different patients (Figure 2C). We observed no sharing of mutations and in fact, very few mutations were found in the stroma samples. This suggests that the stroma in endometriosis lesions is a polyclonal mix of cells and that endometriosis lesions, which comprise both stroma and epithelium, are not seeded by a single ancestor cell giving rise to both cell types. Stroma cells are either recruited to the site by epithelial cells or else stroma cells originating in the normal endometrium travel together with the epithelial cells during retrograde menstruation.

The third observation we make, which holds true for all 14 patients in our cohort with lesional biopsies from multiple body sites, is that epithelial cells isolated from distant body sites have always diverged early in molecular time. An example is shown in Figure 2D. Lesions of superficial endometriosis from the bladder, left pelvis, right adnexa and the right ovary all consists of unrelated cell clones, with no sharing of mutations between cells from different sites. Oligoclonality of individual lesions is further exemplified by the lesion located on the peritoneum of the bladder presenting as rows of cyst-like structures growing in close proximity but with each cyst being clonally derived from an independent cell (Figure 2D-right).

The lack of sharing of mutations between lesions from different body sites suggests that endometriosis does not spread through the pelvis or beyond from an initial founder lesion, similar to metastatic spread of cancer. Rather, each individual lesion is independently seeded and while local clone growth is possible, we find no evidence of clones leaping between sites of the body. That the set of clones found in any two endometriosis lesions from the same patient are always distinct, indicates that endometriosis seeding likely occurs repeatedly throughout life in susceptible individuals.

### Endometriosis cells retain an endometrial identity throughout their lives

Competing theories of endometriosis origin have proposed that lesional cells arise from a non-endometrial origin such as metaplasia from local lineages (*6, 7*) or recruited circulating stem cells (*11, 12*). We tested these cell-of-origin hypotheses using our somatic mutation data from endometriosis lesions. The genomic distribution of somatic mutations along the length of each chromosome has been shown to vary with cell-type specific chromatin organization and epigenetic state (*26*). Thus, somatic mutations preserve a record of the epigenetic landscape experienced by a cell lineage. This principle has been leveraged to infer the cell-of-origin in cancer by correlating distribution of mutations along the genome with chromatin modifications from candidate normal tissue types (*27*).

We applied this framework to endometriosis lesions and eutopic endometrium by constructing mutation profiles and comparing them to epigenomes from 65 normal cell/tissue types. For cells isolated from endometriosis lesions, normal endometrium was the best match (Figure 3; P=6×10^−7^, t-test; Table S4). As expected, cells isolated from the normal endometrium also had a mutational profile most concordant with endometrium epigenomes (P=2×10^−5^), suggesting that eutopic endometrial and ectopic lesional cells share a closely related cell lineage consistent with a common endometrial origin. We also explored the origin of lesion types (superficial, deep infiltrating and endometrioma) separately and again found that normal endometrium was the top-ranked match for each type of endometriosis lesion (P<0.01, Fig. S28).

**Figure 3.**
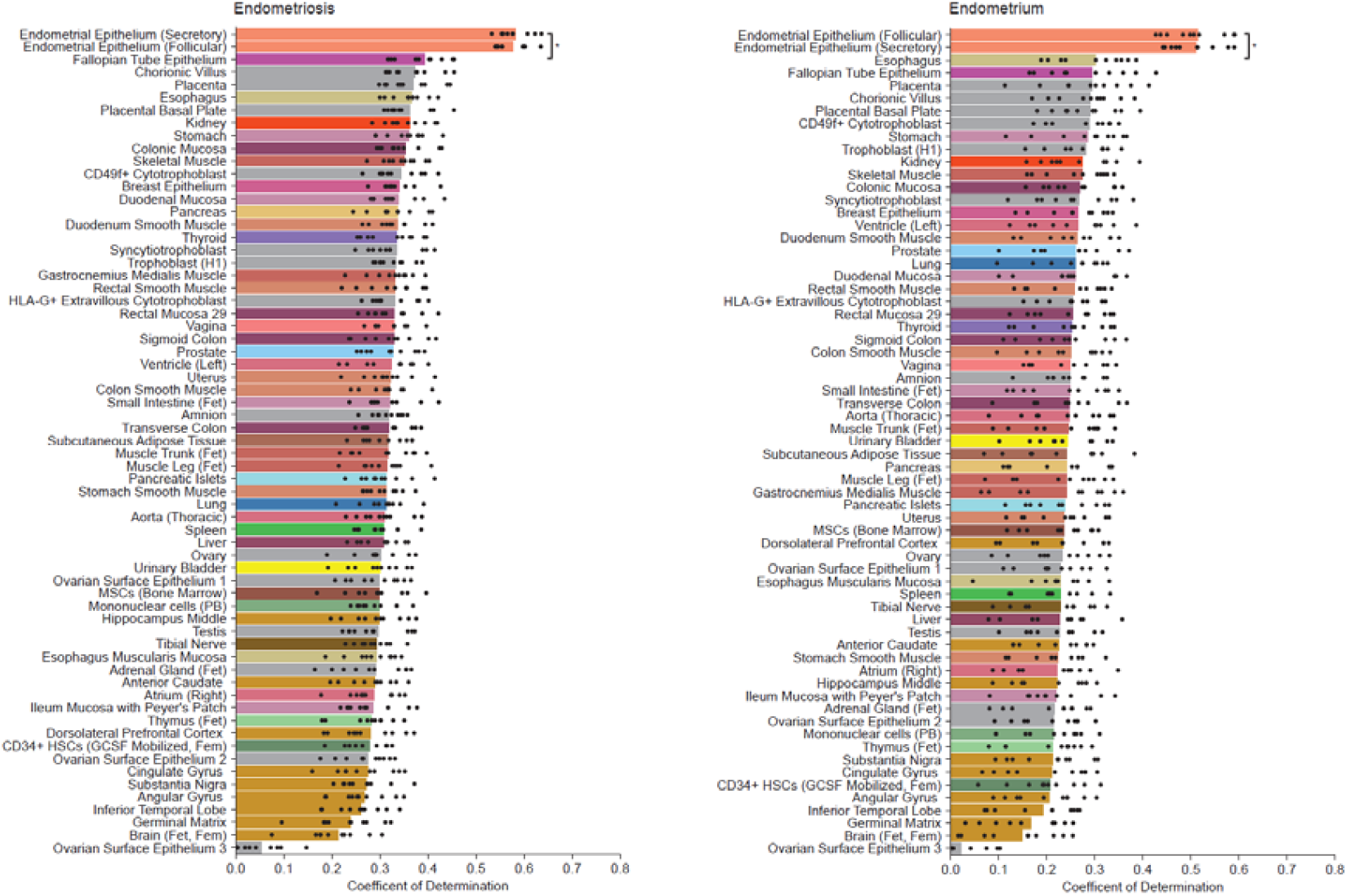
Cell-of-origin analysis for endometriosis (left) and normal endometrium (right). Performance of predicting mutational profiles from histone marks of 65 epigenomes representing distinct cell/tissue types; bars are colored according to tissue category; black points depict values obtained from 10-fold cross-validation; P-values were obtained by comparing the 10-fold cross validation values using a t-test (Abbreviations: HSCs, hematopoietic stem cells; MSCs, mesenchymal stem cells).

We next asked whether a metaplastic transformation could explain lesion development. Under a metaplasia model, mutations acquired after a lineage adopts a new cell identity should reflect the epigenetic state of the transformed cell type, while earlier mutations should retain the ancestral cell type signature. To test this, we classified mutations as early or late depending on their placement on branches of the phylogenetic trees (Methods). When we restricted the analysis to early mutations, normal endometrium remained the best match (P=0.002, Fig. S29), even though we included ovarian surface epithelium epigenome as a proxy for coelomic mesothelium. We also tested the circulating stem cell model by comparing early lesional mutations to hematopoietic and mesenchymal stem cell epigenomes (Fig. S28 and S29), but they showed weaker concordance than endometrium epigenomes, providing no support for a stem cell origin of the epithelial lesions.

Together, these findings consistently support an endometrial lineage origin for endometriosis lesions and argue against a metaplastic origin or a circulating stem cell founder model.

### Large-scale screening of the entire endometrium

The retrograde menstruation theory and theories proposing angiolymphatic spread of endometrial stem cells assume that the cells seeding endometriotic lesions are of uterine origin. We hypothesized that if this is the case, it might sometimes be possible to detect residues of the seeding clones within the normal endometrium. We obtained the entire uteri from three patients undergoing hysterectomies as part of their endometriosis treatments from which lesional biopsies were also available. We used a capture-recapture design to test the hypothesis of uterine origin as follows:

We first used LCM to isolate glands from lesional biopsies for WGS as described above. We built phylogenetic trees from this material and identified independent clones of endometriosis epithelial cells. We defined sets of 10 mutation-barcodes which uniquely identify each clone from the most ancestral branch possible (Figure 4A). We next isolated the entire endometrium from the uteri and divided these up into hundreds of small (approximately 2×2mm) segments, while noting the relative location of each segment (Figure 4B). Lacking positive controls for this assay, we also targeted the *KRAS* G12 and G13 codons and *PIK3CA* codons E542, E545 Q546. Mutations in these oncogenic hotspots are common in endometrial tissue (*15, 16, 28*) but rare in contaminating cell types like blood and stroma. We reasoned that the presence of these mutations in some of the endometrial segments would indicate that our assay was sufficiently sensitive to detect individual mutant endometrial clones given the sequencing depth and the size of the tissue segments.

**Figure 4.**
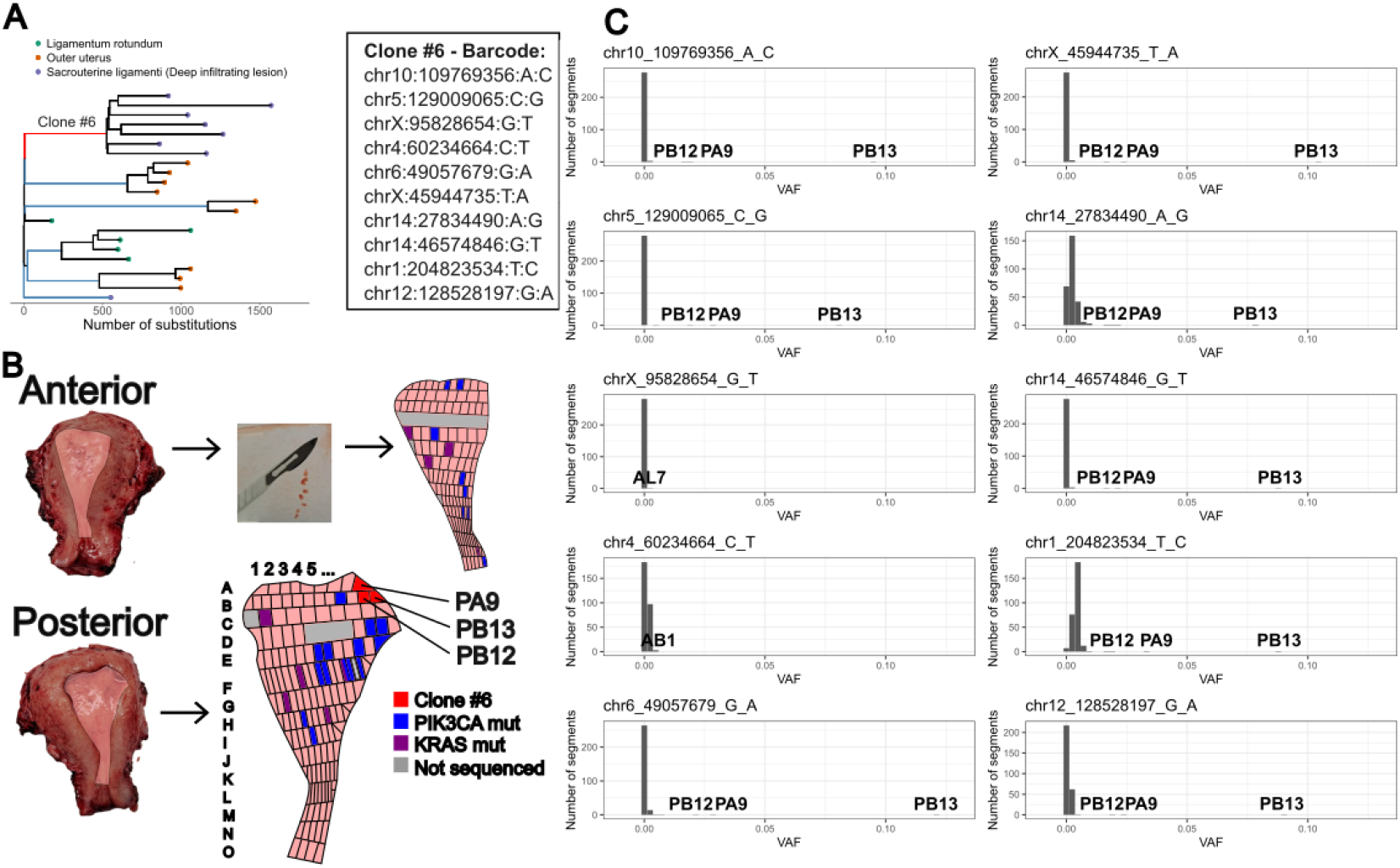
Screening for endometriosis clones in the normal endometrium of patient 42. A) a phylogenetic tree for endometriosis lesions removed during surgery. Branches highlighted in blue and red were selected for the screen. The branch highlighted in red is ancestral to Clone #6, found in a deep-infiltrating endometriosis lesion. B) The normal endometrium of the patient was isolated and dissected yielding 286 segments for the screen. Missense mutations affecting *PIK3CA* E542, E545 or Q546 and *KRAS* G12 and G13 were detected in multiple segments highlighted in blue and purple respectively. Gray denotes parts of endometrium that were retained for clinical work or segments that did not yield sequencing data. Red highlights segments sharing mutations with Clone #6. C) Histograms showing the variant allele frequencies (VAF) of each of the mutations comprising the barcode for Clone #6 in all segments of normal endometrium sequenced. Eight of the ten mutations could be identified in segments PB13, PB12 and PA9.

For each segment of normal endometrium, we used PCR to selectively amplify the regions surrounding the mutations identified in the endometriosis lesions before sequencing, thus achieving a median targeted coverage in the thousands (Methods). As expected, we identified multiple segments carrying missense mutations in the “positive control” *KRAS* and *PIK3CA* hotspot codons in each of the three patients (Figure 4A, Supplementary Text). Furthermore, endometriosis clones could be confidently pinpointed to precise locations of the normal endometrium for two of the patients (Figure 4C and Supplementary Text), revealing those endometriosis lesions to be of uterine origin. Notably, one of the clones traced back to the normal endometrium is from a deep-infiltrating endometriosis lesion dissected from the sacrouterine ligament (Figure 4), showing that seeding of clones from the normal endometrium is not limited to superficial lesions. For the patient highlighted in Figure 4, if 80% of the mutations in the ancestral branch (highlighted in red) occurred before the endometriosis diverged from the normal endometrium, and mutations accumulate linearly with age, then the seeding can be roughly estimated to have occurred when the patient was 17.4-19.2 years old. The results for each patient are further described in the Supplementary Text.

## Discussion

Our data reveal the cellular histories of endometriotic lesions at unprecedented resolution. They confirm that epithelial cells in endometriosis are of uterine origin. Mutational burden, mutational signatures and driver landscapes are highly similar in the normal endometrium as in all types of endometriosis lesions and the distribution of mutations along the chromosomes is reflective of an epigenetic landscape matching the normal endometrium. Furthermore, we isolated cell clones from endometriosis lesions and were able to locate the origin of both superficial and deep-infiltrating endometriosis clones within the normal endometrium. For both clones identified, the high fraction of mutations shared between the lesion and the normal tissue is consistant with lesions diverging from the normal endometrium close to- or after age at menarche.

While our data suggest that cells from the normal endometrium pass into the pelvis via retrograde menstruation, we are unable to determine if this is the only mechanism of spread or whether dissemination is also possible via the angiolymphatic system. However, the presence of multiple unrelated clones within as little as 1 mm of tissue suggests polyclonal seeding of macroscopic segments of endometrium where cells adhere to one another as they travel through the fallopian tube. This is consistent with Sampson’s observation of “bits” of endometrium traveling through the fallopian tubes (*5*) but we cannot exclude a second mechanism involving precise homing of suspended cells to the same location in the body via the angiolymphatic system.

In our screening of entire uteri, we were able to confidently locate 2/14 clones tested within the normal endometrium. In addition to extra-uterine origin, several factors may explain why the remaining clones were not identified. The clones may have migrated entirely to the lesional site without leaving detectable residual cells in the normal endometrium. Alternatively, residual cells in the endometrium may have been outcompeted and replaced by other endometrial clones over time following seeding. It is also possible that these clones resided in tissue segments that were not sequenced, were substantially contaminated with non-epithelial cell types, or were sequenced at insufficient depth to enable reliable detection. The detected clones were isolated from a superficial and a deep-infiltrating endometriosis lesion. Unfortunately, no clones from endometriomas were available for the screen. However, given the similarities of endometriomas to other lesion types at all other levels of analysis, a single causal mechanism is both plausible and parsimonious.

Our results have implications for treatment of endometriosis, in particular because they show that it is likely that the seeding of endometriosis by retrograde menstruation is a repeated event. They support management strategies focused on limiting the number of menstrual cycles to prevent the seeding of new lesions and justify delaying surgery as there is no evidence of existing lesions seeding additional lesions. Oncogenic *KRAS* mutations have been implicated in the pathogenesis of endometriosis (*17*) and several agents targeting *KRAS* are either on the market or under development for treatment of cancers, raising questions about the utility of repurposing such agents for the treatment of endometriosis. We note that although we found *KRAS* mutations in 16% of microdissections, and these often precede clonal expansions within lesions, the mutations are not ancestral to entire lesions, indicating that treatment with KRAS inhibitors would unlikely be curative.

In this study we have provided functional evidence supporting eutopic origin of endometriosis cells as proposed by Sampson in the 1920s. We have further shown that lesions are independently seeded and generally consist of several independent cell clones. It is our hope that understanding of endometriosis etiology will, in time, contribute to the development of treatments and preventions against this often debilitating and understudied disease.

## Acknowledgments

We thank the patients who selflessly donated tissue samples for this study. We further thank the laboratory and informatics teams at Amgen deCODE Genetics for their contribution.

Figure 1A was created in BioRender. Olafsson, S. (2026) https://BioRender.com/d9xryv5.

## Author contributions

SO conceived the project with contributions from KK, AS, VS, IJ, JGJ, ROA and KS. SO and AOA performed microdissections and processed whole uteri with contributions from HSG, BA and GN. HSG performed fixation, sectioning, staining and imaging of tissue samples. SO analyzed the sequencing data and carried out all statistical analyses except for cell-of-origin analyses with contributions from HJ and BA. KK performed cell-of-origin analyses. AS designed PCR primers and oversaw PCR reactions used to amplify regions around mutational barcodes in whole-uteri screening. DM and OM prepared sequencing libraries and oversaw sequencing of all samples. AMJ performed histopathological assessments of tissues and provided guidance for laser capture. ROÁ recruited and consented patients for the study and performed surgeries from which samples were obtained. KS supervised the project. SO wrote the manuscript with contributions from all authors.

## Data and materials availability

The data supporting the findings of this study are available in the supplementary material of this article. Table S1 contains patient-level meta-data. Table S2 contains meta-data and statistics calculated at the level of individual microdissections. Table S3 contains the dN/dScv statistics for each coding gene. Table S4 contains results from the cell-of-origin analysis. Table S5 contains all mutations in *KRAS* and *PIK3CA* identified in the large-scale screen of entire endometrium. Table S6 contains sequences of the PCR primers used in the screen. Read counts for each mutation call, VAF histograms for each sample and tree R-objects will be made available on www.decode.com/summarydata upon article acceptance.

The dNdScv software (RRID:SCR_023123) is freely available at https://github.com/im3sanger/dndscv

The Coselens software (RRID:SCR_022578) is freely available at https://github.com/ggruenhagen3/coselens

The hdp software is freely available at https://github.com/nicolaroberts/hdp

## List of Supplementary Materials

Materials and Methods

Supplementary Text Figs. S1 to S29

Captions for Tables S1 to S6

