## Supplementary Text and figures for "Endometriosis lesions are oligoclonal structures derived from the normal endometrium"

**The PDF file includes:**

Materials and Methods

Supplementary Text

Figs. S1 to S29

Captions for Tables S1 to S6

References

Materials and Methods

1. Sample collection and preparation

1.1 Patient recruitment

Tissue biopsies containing endometriosis lesions or matched normal endometrium were donated by 43 women undergoing laparoscopic surgery for suspected endometriosis at Landspitali University Hospital. Of these, 18 were excluded from the study owing to no- or insufficient amount of endometriosis being detected in the lesional biopsies, leaving 25 participants (denoted P01-P43). All patients gave written informed consent for participation, and the study was reviewed and approved by the Icelandic National Bioethics Committee (ref: VSN-17-188). Unique national identity numbers were encrypted in accordance with regulations imposed by the Icelandic Data Protection Authority.

1.2 Tissue processing and laser capture microdissection

Tissue biopsies suspected to contain endometriosis were fixed in PAXgene Tissue FIX solution (PreAnalytiX, Hombrechtikon, Switzerland) immediately following surgical removal before being transferred to PAXgene STABILIZER solution (PreAnalytiX) for storage. Fixed samples were embedded in paraffin and sectioned. A reference slide for each biopsy was sectioned at 4 µm and reviewed by an expert pathologist. Biopsies found to contain endometriosis were further sectioned at 15-20 µm for laser capture microscopy. Sections were mounted on LEICA polyethylene naphthalate (PEN) membrane glass slides (Leica Microsystems) and stained with hematoxylin and eosin. We used an LMD7 microscope (Leica Microsystems) to dissect individual glands, strips of endometrial tissue and stroma. We combined dissections from serial sections (Z-stacks) whenever possible to increase DNA yield, with a typical sample consisting of 3-6 serial sections. Cells were lysed using the Arcturus PicoPure DNA Extraction Kit (Thermo Fisher Scientific, Waltham, MA, USA) according to the manufacturer’s instructions.

1.3 Processing of whole uteri

Three patients, patient 39, 42 and 43, undergoing hysterectomies donated whole uteri to this study as well as tissue biopsies containing endometriotic lesions. In these cases, we used laser capture microdissection to isolate cells from endometriotic lesions for whole genome sequencing as described above. We next sought to screen the entire endometrium for the presence of mutations found in endometriosis clones. Uteri were placed on ice immediately following surgical removal. Upon receipt, we separated the endometrium from the underlying myometrium and divided it up into hundreds of segments approximately 2x2 mm in size using a scalpel while noting the location of each segment (See figure below). For each patient, a single 2-3 mm thick slice of the endometrium was removed for clinical work and was thus unavailable for this study. The number of segments sequenced was N=192 for patient 39, N=286 for patient 42 and N=274 for patient 43.

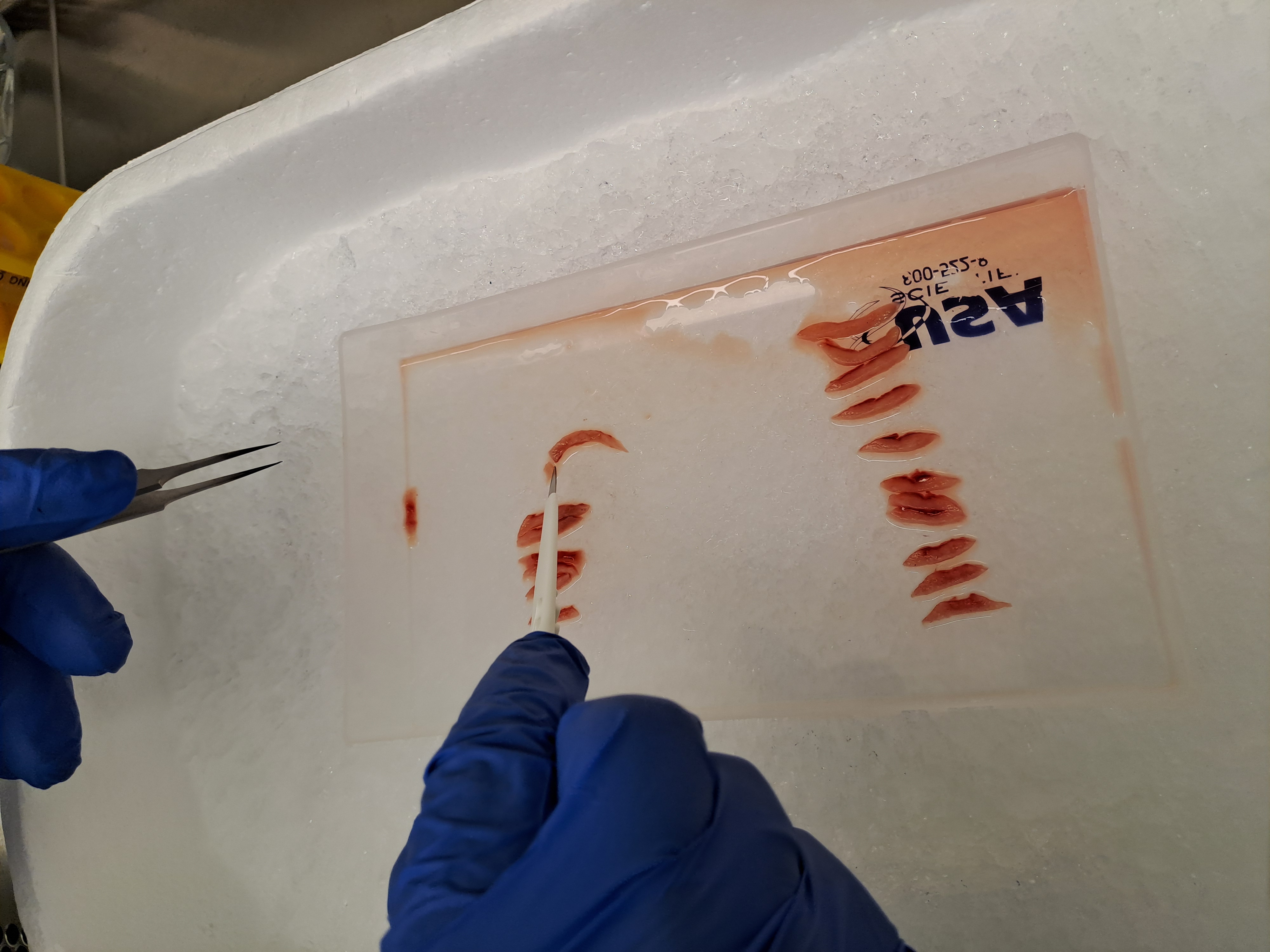

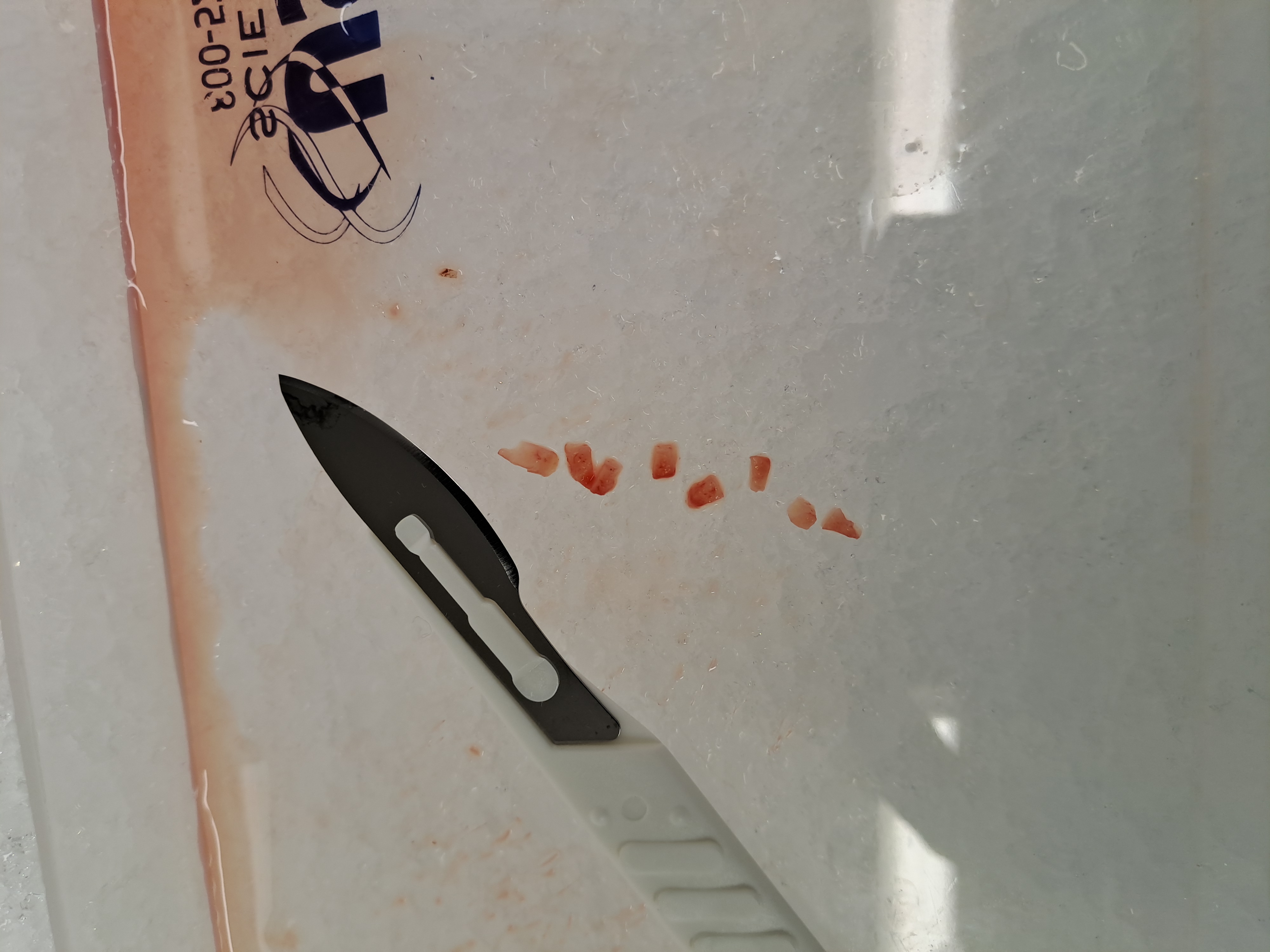

DNA was isolated from each endometrium segment as follows: PBS buffer and a 5mm stainless steel bead (Qiagen) were added to tissue in a 2mL Safe-lock tube. The tissue was partially homogenized in a Tissuelyser LT (Qiagen) for 2 minutes at 50Hz, repeated 3 times. Tubes were centrifuged after homogenization and the PBS discarded. DNA isolation from the tissue was performed using MasterPure Reagents (Biosearch Technologies). In short; Proteinase K digestion at 65°C for one hour followed by overnight incubation at 55°C and RNAseA digestion for 1 hour at 37°C.  DNA isolation was completed following the manufacturer's protocol.

2. DNA sequencing

2.1 Whole-genome sequencing of microdissected samples

To generate sequencing libraries from small inputs of DNA, isolated from a few hundred cells, we followed a previously established protocol (*20*).

Samples were sequenced using 150 bp paired-end reads on Illumina NovaSeq X machines. Owing to the small number of cells sequenced, some samples have low library complexities, which manifests as high fraction of PCR duplicates in the samples. To conserve sequencing resources, all microbiopsies initially underwent low-pass sequencing (target coverage ~1X), based on which samples with high fractions of PCR duplicates and/or near-zero coverage were excluded. Remaining samples underwent a second round of sequencing to achieve higher depth. Some samples had >5X coverage after the first round of sequencing but were nevertheless not selected for deeper sequencing. However, we have included such samples in our analysis but note that sensitivity for mutation calling for samples with such low coverage will be limited. This partially explains why some branches of the phylogenetic trees appear shorter than the rest. For the cohort as a whole, the average sequencing coverage of individual microdissections was 18.9X (range, 5-106X, first and third quartiles 14.1X and 25.2X, respectively). Sample sizes quoted in the main text refer to samples with >5X coverage.

Following demultiplexing, reads were aligned to the hg38 human reference using bwa (*29*) (v 0.7.17). Duplicate reads were marked and coverage and duplicate statistics were calculated using Picard (v 2.20.3) (<http://broadinstitute.github.io/picard>).

2.2 Targeted sequencing of whole uteri

We screened hundreds of segments of endometrium from three uteri for the presence of somatic mutations found in endometriotic lesions from the same donors. In total, we screened for 140 mutations tagging 14 endometriosis clones: 2 clones from patient 39, 6 clones from patient 42 and 6 clones from patient 43. We selected mutations from the most ancestral branches of the phylogenetic trees for inclusion in the screen as follows:

1. The mutation should be non-coding substitutions (not indels) and be unlikely to be a driver mutation, as these have a higher probability of being recurrent, and thus not necessarily indicative of shared ancestry.
2. We should be highly confident that the mutation is assigned to the correct branch of the phylogenetic tree.
3. We require at least one of the microdissections in which the mutation is found to have at least seven reads supporting the mutation. Two or more of these reads should map to either strand of the DNA molecule.
4. The mutation should not map to any of the following:
   a. Centromeric regions.
   b. A list of putative mutations/artifacts called in a panel of Icelandic normal samples. An Icelandic panel of normals was constructed by running CaveMan on 100 samples of whole-blood from individuals younger than 35 years that have participated in past genetic studies at deCODE, using the sample from another randomly selected individual as unmatched normal. We reasoned that these individuals are extremely unlikely to have clonal haematopoiesis and any mutation called would most likely represent a recurrent artifact or a germline variant not captured by the CaveMan default filters (which include comparison to a panel of normal generated at the Wellcome Sanger institute (*30*)).
   c. Regions containing putative germline variants in the deCODE database (daisy freeze) but which fail quality control.
   d. Microsatellite polymorphisms with over 15 alleles in the Icelandic population (*31*).
   e. A genomic mask designed to capture low-quality sites in the Gnomad database and sites with high error rates at the Wellcome Sanger institute (*32*).
5. From mutations passing the above criteria, we randomly selected 15 mutations from each clone for manual inspection in a genome browser and from that set, selected 10 to make up the barcode for the clone.

There is no positive control for this screen and if no mutations were observed, we would not know if this was due to the assay being insufficiently sensitive to detect individual clones in our endometrial segments. To increase our confidence that we could observe endometrial clones using this design, we included in the screen hg38:chr12:25245346-25245351 and hg38: chr3:179218294-179218307, the sites encoding G12 and G13 of KRAS and E542, E545 and Q546 of PIK3CA. Hotspot mutations affecting these codons are commonly observed in normal endometrial epithelium (*15, 17*) but are not be expected at high frequencies in contaminating cell types such as stroma, myometrium or blood. The presence of these mutations would indicate that we detect endometrial clones and provide an estimate of the minimum VAFs of the clones that could be detected. We acknowledge as limitations of this strategy however, that clones carrying strong driver mutations may be larger than average and it is conceivable that more than one clone, each carrying an independently acquired hotspot mutation, may be present in the same endometrial segment. In both cases, inflated VAFs would be expected.

To perform ultra high-depth sequencing, we first used PCR to amplify across the sites of interest across all endometrial segments. Including the positive controls, 22, 62 and 62 sites were amplified for patients 39, 42 and 43, respectively (for a total of $22\times191+62\times286+62\times274=38,922$ reactions). For each site, two sets of primers were designed using the Primer 3 software ([www.broadinstitute.org](http://www.broadinstitute.org/)). Primer sequences are available in Table S6. PCR reactions were set up on Zymark SciClone ALH300 using Taq polymerase (Fermentas) and run on MJ Research PTC-225 thermal cyclers. All PCR products for each segment were pooled into one well and purified using AMPure XP (Agencourt) in a 1:1 ratio. Purified samples were prepared for Illumina sequencing using the NEBNext Ultra™ II PCR-free kit from New England Biolabs. Pooled amplicons were fragmented in 96-well TPX-AFA plates (Covaris Inc.) using the Covaris LE220plus instrument with the following settings: sample volume = 50 µL, duty factor = 25%, cycles per burst = 50, temperature 10 °C, peak incident power = 200W, treatment time = 250 sec. End repair and A-tailing was performed in a single step followed by ligation of unique dual indexed sequencing adaptors (IDT for Illumina) and two rounds of SPRI-bead (Beckman Coulter) purification (1:1) using the Hamilton STAR liquid handler. Sequencing libraries were quantified by Qubit (dsDNA), pooled in equimolar amounts and sequenced on the NovaSeq X sequencer on a single lane of the 25B flowcell (Illumina). Sequencing was done using the paired-end workflow with a read length of 2*150 cycles of incorporation and imaging in addition to 2*8 index cycles for the molecular barcodes.

3. Mutation calling in microdissected WGS samples

Point mutations were called in four steps as previously described (*21*): Initial discovery, initial filtering of the discovery set, genotyping and final filtering. For initial discovery of indels and substitutions, we used cgpPindel (*33*) and CaveMan (*30*), respectively. WGS data from blood was available for 6/25 donors who have been sequenced as part of past research projects at deCODE (*34*). When WGS from blood was available, we used that as a matched normal sample. For two other patients, we used a polyclonal matched normal sample comprising a mix of muscle, non-endometriosis stroma, lymphocytes and/or connective tissue from different sites of the biopsy and isolated with LCM. For the remaining 18 patients, we called mutations using an unmatched normal from a different patient.

In the initial filtering step, we removed variants that did not pass default filters for Pindel/CaveMan. For substitutions, we further filtered mutations if the medial alignment score of reads reporting the alternative allele was <140.

In the genotyping step, we compiled for each patient a list of putative mutations found in any sample from that patient. We next counted the number of reads reporting the reference and alternative alleles for each mutation in each sample using the bam2R() function of the deepSNV package (*35*). Only properly paired reads with mapping quality >30 and bases with Phred scores >30 were counted and reads matching the SAM flag 3844 were excluded (reads that are unmapped, not primary alignment, fail platform/vendor QC, are supplementary alignments or reads that are PCR or optical duplicates).

To filter the mutations thus genotyped, we next applied a set of binomial filters designed to capture unfiltered germline variants and recurrent sequencing artifacts (*21*). Germline variants will be present at a high VAF in all samples from an individual. We tested if the read counts from the previous genotyping step were consistent with the reads being drawn from a binomial distribution with a probability of success of 0.5. The alternative hypothesis is that reads are drawn from a distribution with a lower probability of success. We calculated P-values for all putative mutations found in a patient, performed Benjamini-Hochberg correction for multiple testing and excluded mutations where we failed to reject the null hypothesis (q>0.01). Recurrent sequencing artifacts are often found on a few of reads across many or all samples. Because they are not found on half the reads, recurrent artifacts pass the exact binomial test described above. While recurrent sequencing artifacts are present at a low level across multiple samples, true somatic mutations should have multiple supporting reads in one or a few samples and be entirely absent in others. In other words, they have a higher overdispersion than recurrent artifacts. We fitted a negative binomial model to the read count matrices to estimate $\boldsymbol{\rho}$, the overdispersion factor of the model. We excluded mutation calls with $\boldsymbol{\rho}$<0.1 as in our previous work (*21*).

A final filtering step involved removing mutations that could not be unambiguously mapped to the phylogenetic trees, as described below.

4. Building phylogenetic trees

Phylogenetic trees were constructed for each patient using the MPBoot software (*36)* (v. 1.1.0) as previously described (*21*). Briefly, MPBoot constructs a maximum parsimony consensus tree for each patient using ultrafast bootstrap approximation. Mutations were then assigned to the branches of the tree using a maximum likelihood approach (*21*). Since substitutions are generally called with higher confidence than indels, we generated the trees based on the substitutions only and then mapped indels to the branches of those trees. We used a likelihood ratio test to compare the likelihoods of the mutation belonging to the assigned branch versus any other branch of the tree and filtered mutations with p>0.05 of belonging to a different branch than the one assigned.

The phylogenetic trees presented in Figure 2 and Figures S4-S27 sometimes show significant variance in branch length. This is in large part due to technical variance. We included in the trees microdissected samples sequenced to as low as 5X coverage (Figure S1). As shown in Figure S2 below, sensitivity of the mutation calling pipeline is drops at low coverage.

5. Estimating sensitivity of mutation calls in microdissected samples

For two normal endometrial glands, we dissected adjacent histological sections of the same gland twice and made independent sequencing libraries for each pair. Sequencing libraries were independently created and samples were sequenced to >30X coverage and then down-sampled to estimate the sensitivity of our pipeline at different sequencing coverages. Sensitivity at coverage X, S_X_, was estimated as:

$$S_{x}=\frac{2\times n_{2,X}}{n_{1,X}+2\times n_{2,X}}$$

Where n_1,X_ is the sum of the number of mutations found in just one of the members of a pair at coverage X and n_2,X_ is the number of mutations found in both members of a pair at coverage X.

The results from this analysis are shown in Figure S2a below. The figure shows that sensitivity is ~75% at 10X coverage, rising to ~88% at 20X and >90% at 30X.

We did not formally quantify the specificity of our pipeline. However, as shown in Figure S2b-c below, few mutations were unique to members of each pair, indicating high specificity.

6. Mutation burden estimation

To compare the mutation burdens of endometrial cells isolated from lesional and normal biopsies while controlling for non-independent sampling, coverage and median VAF, we used linear mixed-effect models as implemented in the nlme (v. 3.1-166) package in R as previously described (*21*). Briefly, we used the total number of mutations detected as a response variable and included in the model fixed effects for year of birth in 5-year bins, the coverage and the median variant allele fraction of each microdissection. Observations are non-independent and we included in the models random slopes for patient and for biopsy. The biopsy effect was nested within that for the patient. As many early developmental mutations will be filtered as germline, we constrained the intercept to zero, the biological interpretation being that no somatic mutations are present at birth. To allow for differences in variance in mutation burden between the normal endometrium and endometriosis from different body sites, we constructed a general positive-definite variance-covariance matrix for the random effects of patient and biopsy by sample type. We fitted models with and without a fixed effect for lesional/normal status and used likelihood ratio test to test if including this variable significantly improved the fit of the model.

While endometrial glands represent clonal populations of cells, clonal boundaries are not visible for endometriomas, where endometrial cells are often organized into thin strips of tissue as opposed to defined glands. In these cases, individual microdissections may comprise more than one cell clone, which would manifest as skewing of the VAF distribution from 0.5 toward 0. We repeated our modelling using only samples with median VAF>0.3 but found no differences in the mutation burden between normal and lesional samples.

7. Mutational signature extraction

We used the hdp package (https://github.com/nicolaroberts/hdp) in R (v. 0.1.5) to perform mutational signature extraction. To avoid double-counting, mutations were mapped to branches of the phylogenetic trees and each branch over 50 mutations in length was treated as a single sample by the signature extraction algorithm. The signature exposures for each branch were then weighted by the branch length to create an exposure for each microdissection sample.

An advantage of the hdp approach is that it can both fit against know signatures and discover novel signatures simultaneously. To facilitate the detection of previously identified signatures known to be commonly found across a range of normal tissues (*13*) and in normal endometrium in particular (*15)*, we conditioned hdp with the following single-base substitution signatures: SBS1, SBS2, SBS5, SBS8, SBS13, SBS18. The conditioning was done by assigning to frozen nodes 20,000 pseudocounts. During the Dirichlet process, the frozen pseudocounts are unable to leave their initial clusters but counts from data nodes may join these clusters.

The hyperparameters for the α clustering parameter were set to 1. The model was initiated with 7 clusters (one more than the number of prior signatures). After discarding as burn-in the first 100,000 iterations of the Gibbs sampler, we sampled 100 posterior samples at intervals of 2,000 iterations. We combined for signature extraction the results of 20 such chains. We merged clusters with cosine similarities >0.85.

In addition to the null cluster, seven mutation clusters (hdp-components) were identified by the algorithm (Figure S3). Four corresponded to COSMIC signatures; SBS5, SBS18, SBS2, and SBS13. Three “novel” components, hdp-N1-3 did not match any COSMIC signature. The first of these, hdp-N1, accounted for a large fraction of the mutations in the cohort and was clearly a mix of SBS1, SBS5 and SBS18, which are ubiquitous in the normal endometrium (*15*) and in many other normal tissues (*13*) and were likely too correlated within samples in our cohort to be separated by the algorithm. In this case, we used expectation maximization to deconvolute the hdp-N1 component into SBS1 (28%), SBS5 (54%) and SBS18 (18%) as previously described (*37*). We reconstructed the component using these signatures and found the reconstructed component to have cosine similarity of 0.99 with hdp-N1.

The second “novel” hdp component, N2, is a flat signature which shows high cosine similarity with SBS5 and is a close match with the signature of endogenous mutations found in blood (*38*) (cosine similarity 0.96). We hypothesize that this is a manifestation of similar cell-endogenous mutational processes as those thought to give rise to SBS5 and may reflect tissue-to-tissue variability in SBS5. We merged hdp-N2 with SBS5 in all subsequent analyses.

The third “novel” hdp component hdp-N3 was characterized by C>A mutations and found at a low level across all samples. We hypothesize that this component may represent sequencing errors which remain in our dataset after filtering. Oxidating damage to sequencing libraries, and low-input libraries in particular, is known to result in C>A artifacts (*20*). We therefore combined the hdp-N3 component with the “Unassigned” component.

8. Positive selection and driver assignment

We used dNdScv (*22*) (v. 0.0.1.0) to detect genes under positive selection in the dataset. To avoid counting single mutation events multiple times, we conservatively treated each branch of the phylogenetic trees as a sample in this analysis. Combining all mutations in the cohort revealed enrichments of mutations in *PIK3CA, KRAS; FBXW7, FIK3R1, ARHGAP35* and *FOXA2* (see main text), which survived Benjamini-Hochberg correction for multiple testing. We repeated the analysis after stratifying mutations depending on whether they were found in endometriosis or normal endometrial biopsies. In each case, no additional genes were identified and only *KRAS* and *PIK3CA* reached significance in either cohort (Table S3).

To formally test for differences in positive selection between endometriosis and normal endometrium, we used the Coselens software (*23*) (v.1.0). Coselns uses the dN/dS ratios estimated by dNdScv to estimate the number of drivers per sample and compares the number of drivers in two groups using a likelihood ratio test (*23*). It was developed for cancer sequencing studies that sequence a single sample per individual. Treating each branch of the phylogenetic tree as a sample skews the Coselens estimates when different numbers of coalescent events are observed for each group, either due to unequal sampling or differences in clonal composition. We therefore randomly assigned mutations detected in endometriosis and normal endometrium to pseudosamples of 1,000 mutations each. This changes the interpretation of the effect estimates from Coselens. Rather than estimating a difference in the number of drivers per sample (tumour), we now estimate an average difference in the number of drivers per 1,000 mutations.

**9. Cell-of-origin analyses**

The distribution of mutations along the length of chromosomes is a record of the epigenetic state of a cell and this can be used to infer the tumor cell-of-origin based on the somatic mutations alone (*27*). Following prior work (*27*), we binned the genome into 2,128 windows of 1 Mb spanning ~2.1 Gb of DNA and modeled the somatic mutation density from chromatin immunoprecipitation-sequencing (ChIP-seq) reads using random forest regression. Models were evaluated in a 10-fold cross-validation setting using the LogCosh distance between observed and predicted mutation profiles. We determined the difference between models on the values from the tenfold cross-validation using a corrected t-test for cross-validation error estimates (*39*).

After excluding mutations detected in adenomyosis or cesarean section scar tissue, we assigned each mutation to a tissue compartment (normal endometrium versus endometriosis lesions) and aggregated mutations within each compartment to generate mutational profiles. Similarly, mutations from the lesional biopsies were grouped and aggregated according to lesional type (i.e., superficial endometriosis, deep-infiltrating endometriosis and endometriomas). We also stratified and aggregated mutations into early- versus late-acquired events when possible. The underlying hypothesis here is that a non-endometrium cell-of-origin could be masked by the accumulation of mutations occurring after cell-conversion. To enrich for early occurring mutations, we assigned mutations as late-occurring if they followed a coalescent event that occurred after >20% of the molecular age of the clade-node as determined by the highest mutation burden of its children.

Epigenetic data, including the histone marks H3K27me3, H3K9me3, H3K36me3, H3K4me1, H3K27ac and H3K4me3 together with the matched input control track, were obtained from Roadmap (*40*), ENCODE (*41*) and IHEC (*42*), supplemented with published datasets (*43*). In total, we used the top three histone marks of 65 epigenomes from distinct tissue/cell types for the predictions.

**10. Mutation calling from deep sequencing of whole uteri**

We hypothesized that if endometriosis cells originate in the normal endometrium, it might in some cases be possible to detect cells belonging to the ancestral clone still residing in the normal endometrium. Section 1.3 above describes how entire uteri from three patients were dissected into hundreds of segments. We constructed barcodes of 10 mutations each to tag clones identified in endometriosis biopsies and performed PCR amplification and targeted sequencing to identify the mutations in the endometrial segments as described in section 2.2 above. What follows is a description of the mutation calling from this material.

The sequencing of the PCR amplicons resulted in extremely high and often variable sequencing coverage across sites and segments. The depth quantiles for each patient are:

|  | 0% | 25% | 50% | 75% | 100% |
| --- | --- | --- | --- | --- | --- |
| Patient 39 | 0 | 91,603 | 180,725 | 339,952 | 983,601 |
| Patient 42 | 0 | 6,963 | 21,972 | 55,910 | 1,943,044 |
| Patient 43 | 0 | 25,332 | 77,981 | 161,147 | 1,616,129 |

Note that these numbers represent depth without removing reads marked as duplicates. The extremely high depth achieved over relatively short amplicons means that we are likely to observe by chance multiple reads having identical start and end positions after mapping that may or may not derive from the same fragment. We performed calling as described below with and without removing reads marked as duplicates and observed comparable results.

To call somatic mutations we used the ShearwaterML algorithm, as implemented in the deepSNV package (v. 1.50.0) in R. ShearwaterML was designed to detect mutations present at a low variant allele fraction in samples sequenced to a high depth (*35, 44*). Briefly, the algorithm constructs a compound reference by aggregating data across all segments from the same individual. The error rate at each base is modeled by fitting a beta-binomial model to the compound reference and for each segment we test if the number of reads reporting the alternative allele is greater than expected given the depth and the base-specific error model. If i is a sample index, j is a site index and $k\in\{A,C, G, T,-\}$, where ‘-‘ represents the deletion of a base, then the number of reads reporting each allele on the forward and the reverse strands ($X_{ijk}$ and ${X'}_{ijk},$ respectively) can be modeled as being drawn from a beta-binomial distribution:

$$X_{ijk}\sim BetaBin\left( n_{ij},v_{ijk},\rho_{jk} \right)$$

$${X'}_{ijk}\sim BetaBin({n^{'}}_{ij},{v'}_{ijk},\rho_{jk})$$

Where the parameters $n_{ij}$ and ${n^{'}}_{ij}$ represent the total coverage at the site in the sample, and the parameters $v_{ijk}$ and ${v'}_{ijk}$ represent the mean fraction of reads reporting a given base across all samples in the cohort. In other words, $v_{ijk}$ and ${v'}_{ijk}$ represent the average error rate of a given base at a given site, assuming that most segments will be wild-type for a given mutation at a given site. Finally, $\rho_{jk}$ represents the overdispersion factor of the error rate at the site, i.e how much this varies between samples in the set. We used the betabinLRT() function of the deepSNV package to fit the model, setting the rho parameter to NULL such that it is estimated from the data.

The null hypothesis is that the number of reads from each strand reporting the alternative allele is drawn from the beta-binomial distributions $X_{ijk}$ ${X'}_{ijk}$. The alternative model in contrast assumes reads are drawn from beta-binomial distributions with mean $\mu_{jk}=\mu_{jk}^{'}$:

$$H_{0}=\mu_{jk}=v_{jk},\mu_{jk}^{'}=v_{jk}^{'}$$

$$H_{1}:\mu_{jk}\left( {=\mu}_{jk}^{'} \right), v_{jk},v_{jk}^{'}$$

For each site, the likelihood of both models is calculated for each possible nucleotide different from the reference. A P-value for each mutation call is calculated from a likelihood ratio test with two degrees of freedom on the likelihood of both models. We adjusted the outputted P-values of mutation calls using Benjamini-Hochberg correction for multiple testing. We also filtered calls if the coverage of a site in a particular segment was less than 200X. We further required at least 5 reads mapping to either strand to call a mutation.

Supplementary Text

Screening of the whole endometrium

We received uteri from hysterectomies from five of the 43 participants. Of these, matched endometriosis lesions were available for three participants, Patient 39, Patient 42 and Patient 43, which we focus on below.

Patient39

Patient 39 was the first patient screened and is also the youngest participant in the study (born in 1992-1997). The uterus was cut open and a section from the topmost part of the anterior part was removed for clinical assessment.

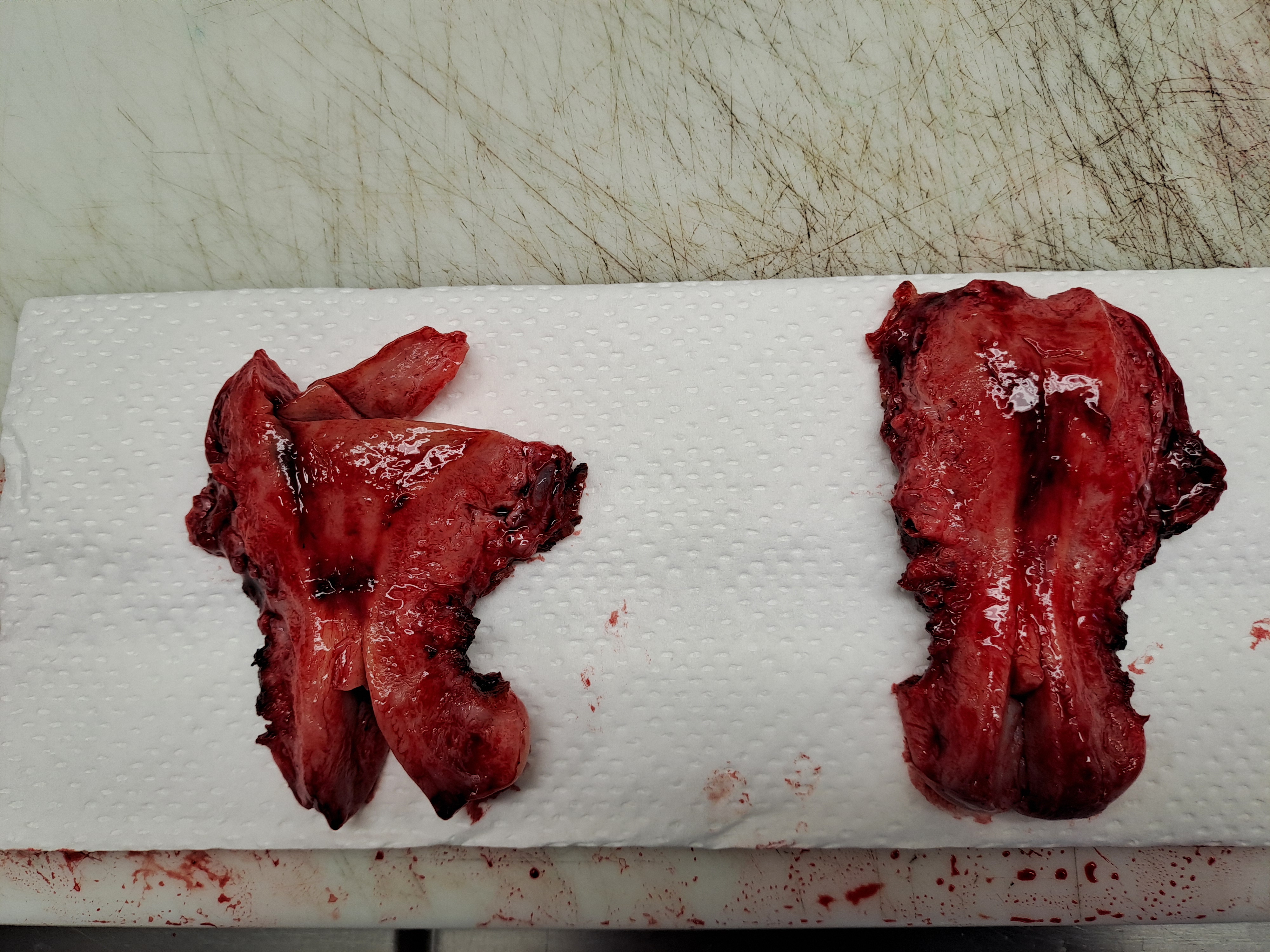

**The uterus donated by Patient 39**

Patient 39 presented with superficial endometriosis. Two biopsies of endometriosis lesions were available for the study, one from the left pelvis and one from the peritoneum of the bladder. Following sectioning and H&E staining, multiple endometriosis glands were visible from the biopsy from the left pelvis. These were dissected and later found to all belong to a single clone (See panel A of the figure below). This was labeled Clone #1. From the biopsy taken from the bladder peritoneum, only a single gland could be isolated and sequenced and this single sample comprises Clone #2 for this patient.

We selected mutations as barcodes, performed PCR and sequencing as described above. Two 96-well plates, containing 191 sample were initially sequenced, revealing that 5/10 mutations comprising Clone #1 could be identified in segment AC8 (panel B below). Having identified the clone, we refrained from sequencing some remaining segments isolated from the posterior part of the uterus (marked in gray in the figure).

Clone #2 from Patient 39 could not be identified in the segments sequenced. We note that the mutations selected for this clone are subject to additional limitations compared with the barcode mutations for Clone #1; namely 1) The mutations were only found in a single sample and so are more likely to represent sequencing errors or artifacts, while the barcode for Clone #1 consists of mutations independently identified across multiple samples and sequencing libraries. 2) The barcode for Clone #2 is less enriched with early mutations. Since there is no coalescent event for this clone to split mutations into early vs late occurring, a larger fraction of the mutations selected for the barcode may have occurred post-seeding in Clone #2 compared with Clone #1 and would thus not be found in the normal endometrium.

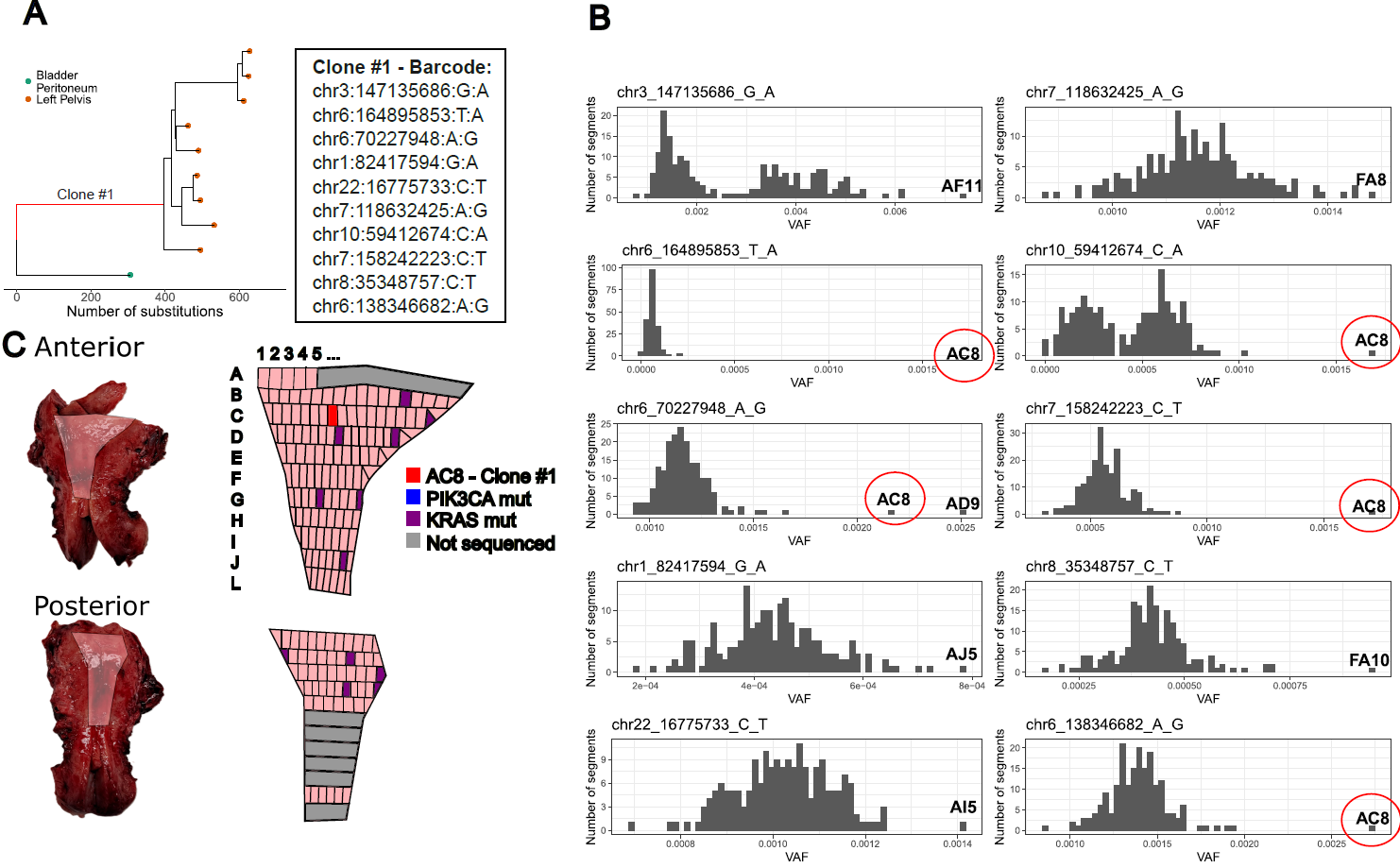

**Screening for endometriosis clones in the normal endometrium of patient 39.** Two endometriosis clones were selected for the screen. Results are shown for Clone #1. A) a 10 mutation barcode was selected from the most ancestral branch possible. B) The whole endometrium was isolated and divided into hundreds of small (approximately 2x2mm) segments. The top gray part of the anterior half was removed for clinical work and was unavailable for the study. The gray areas of the posterior half represent segments that were isolated but we refrained from sequencing because the clone in AC8 had already been identified in the first 191 segments sequenced. C) Histograms showing the variant allele fraction (VAF) of each of the ten mutations comprising the barcode for Clone #1 and highlighting the endometrial segment with the highest VAF. Five of the ten mutations are called in segment C-8 from the anterior part of the endometrium, circled in red. This segment shows the highest VAF for all five mutations except for one, where the highest VAF is in AD9, a segment directly adjacent to AC8.

The time of divergence of the endometriosis clone from the normal endometrium can be roughly approximated as follows: Since five of the ten barcode mutations were shared, we estimate that 50% of the mutations in the ancestral branch occurred before divergence. Dividing this number by the average total mutation burden of each of the leaf-nodes of the clade gives the fraction of mutations which occurred before the divergence. Assuming a linear accumulation of mutations with time, we estimate that the seeding of the endometriosis clone occurred when the patient was 10.8-12.7 years old. This is younger than the self-reported age at menarche for this patient, which is 15 years. However, we caution against over interpreting this as the 50% estimate is subject to uncertainty owing to the small number of barcode mutations assessed. If only one or two more mutations could have been identified in the endometrium, then the estimated age at seeding would be consistent with the age at menarche.

Patient42

We next processed a uterus donated by Patient 42. Upon inspection of the uterus, we noted two putative endometriosis lesions on the outside of the uterus. These were first dissected and embedded in paraffin. Upon pathological assessment, one was found to contain endometriosis and was included in the study, as well as two further endometriosis lesions removed during the surgery; One superficial lesion from the ligamentum rotundum and one lesion burrowed >5 mm within the sacrouterine ligament, which is a deep-infiltrating lesion.

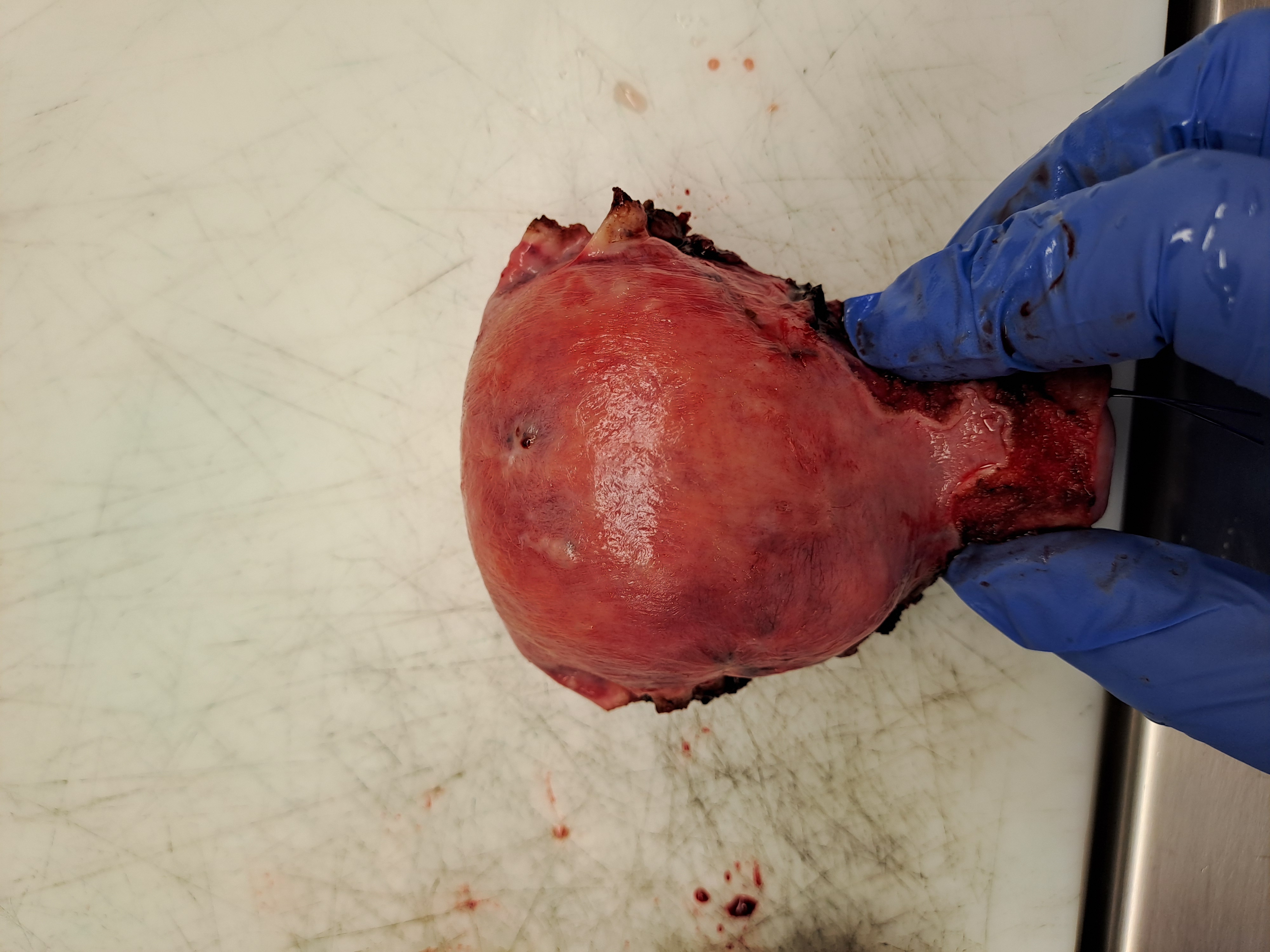

**The uterus donated by Patient 42. Encircled in yellow are two putative endometriosis lesions removed before sectioning of the endometrium.**

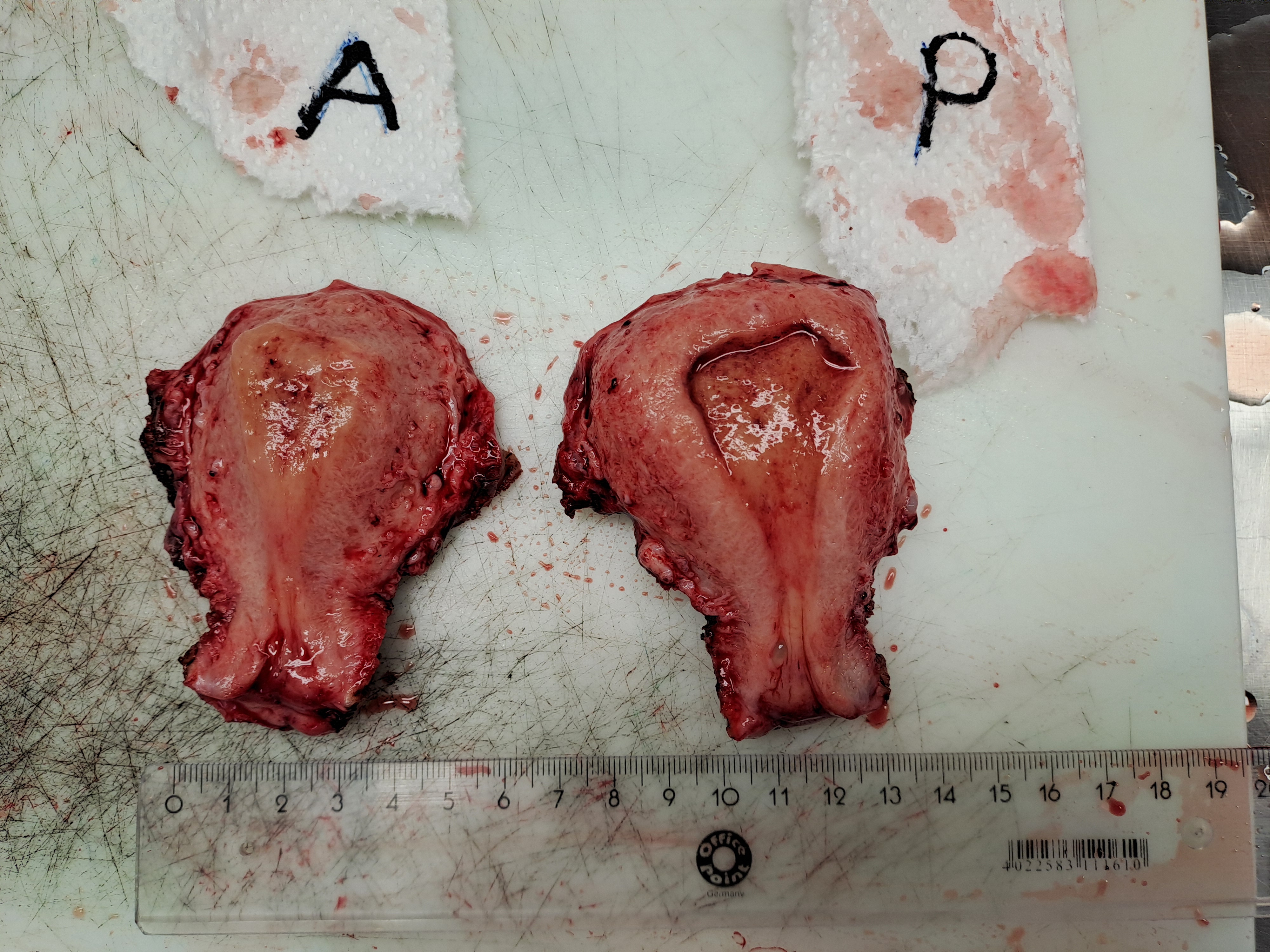

**The uterus donated by Patient 42 after opening.**

A slice of the endometrium from the anterior and a middle part of a slice from the posterior were removed for clinical work and were unavailable for the study (marked in gray in the figure below).

From three endometriosis biopsies, we sequenced 21 individual glands and constructed phylogenetic trees as described above. We selected barcodes tagging six clones in the tree. The results are described in Figure 4 of the main text which we re-publish here for ease of reading. Mutations from one of the clones, Clone #6, comprising the glands isolated from the deep-infiltrating lesion of the sacrouterine ligament, was clearly visible in three adjacent segments of normal endometrium. In one segment, PB13, the mutations were found at 8-12% variant-allele frequencies, meaning that 16-24% of all cells in the segment carried the mutation. This clone had clearly expanded to a macroscopic scale within the normal endometrium prior to seeding of the endometriosis lesion.

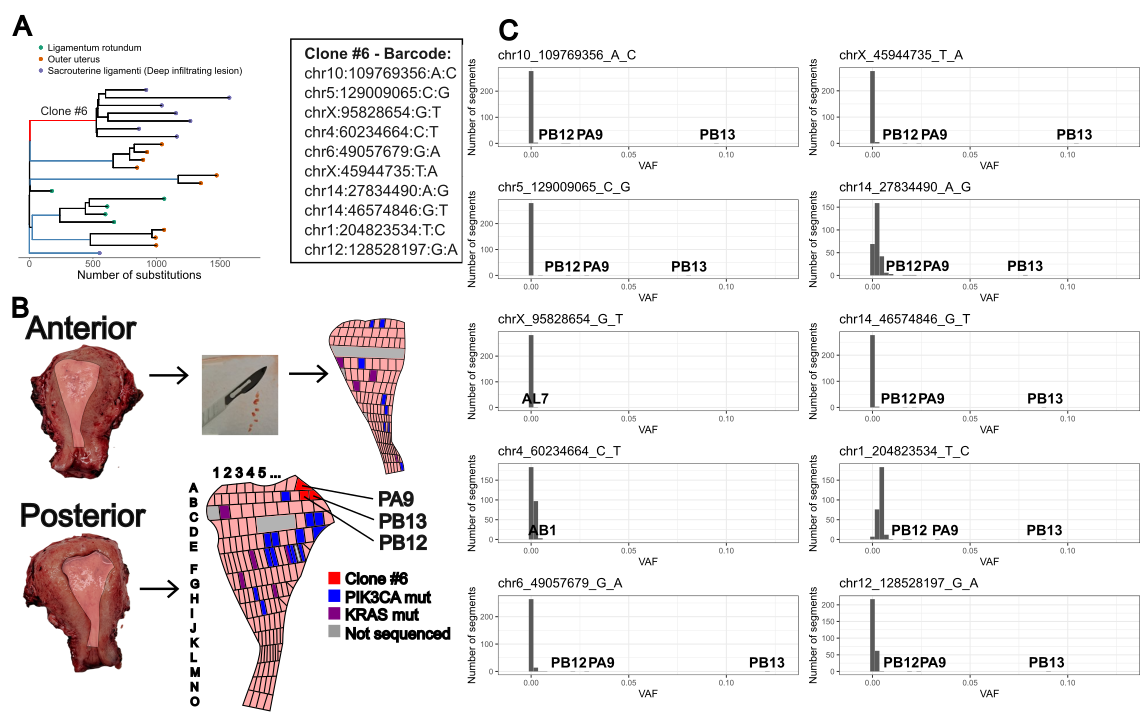
 **Screening for endometriosis clones in the normal endometrium of patient 42**. A) a phylogenetic tree for endometriosis lesions removed during surgery. Branches highlighted in blue and red were selected for the screen. Clone #6 is ancestral to a deep-infiltrating endometriosis lesion and is further highlighted in red. B) The normal endometrium of the patient was isolated and dissected yielding 286 segments. Missense mutations affecting PIK3CA E542, E545 or Q546 and KRAS G12 and G13 were detected in multiple segments highlighted in blue and purple respectively. Gray denotes parts of endometrium that were retained for clinical work or segments that did not yield sequencing data. Red highlights segments sharing mutations with Clone #6. C) Histograms showing the variant allele frequencies of each of the mutations comprising the barcode for Clone #6 in all segments of normal endometrium sequenced. Eight of the ten mutations could be identified in segments PB13, PB12 and PA9.

Following a similar process as for Patient 39 above, dividing the length of the branch by the average mutation burden of all children nodes and multiplying by 0.8, we estimate that the endometriosis diverged from the normal endometrium when this patient was 17.4-19.2 years old. This is after the self-reported age at menarche, which is 13 years old. It is worth noting again that the outcome of this exercise depends on the fraction of mutations shared between the ancestral branch of the clone and the normal endometrium, which is estimated using very few mutations and thus is subject to uncertainty.

Patient43

Finally, patient 43 donated a uterus to the study.

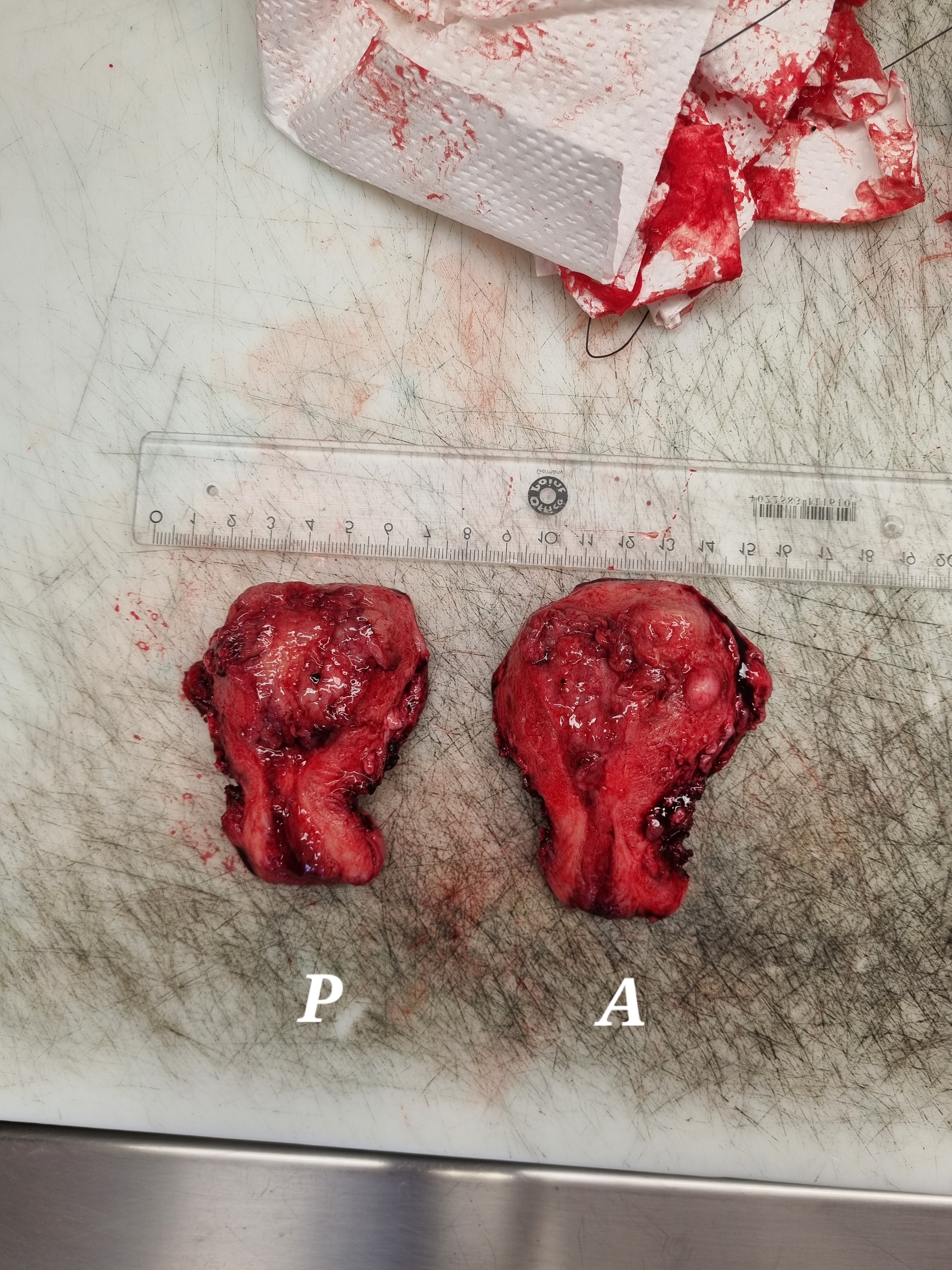

**The uterus donated by Patient 43.**

The patient presented with superficial endometriosis and biopsies were available from lesions on the left ovary. We sequenced 24 endometrial glands from the endometriosis lesions and constructed phylogenetic trees as described above. The 24 glands comprised 13 independent clones (panel A in the figure below). We chose 6 clones for inclusion in the screen where the clones comprised two or more samples. As discussed for patient 39 above, the inclusion of clones comprising multiple samples has the added benefit of the mutations having been independently identified in two or more samples, and it is possible to enrich for ancestral mutations by selecting mutations that map to branches preceding coalescent events. However, we note that due to the high degree of sharing (recent split) between samples in many of the clones from Patient 43, and small sizes of the clones (no clone comprises more than 3 glands), the screening of the mutations in this patient is subject to some of the same limitations as noted above for Clone #1 for patient 39, albeit to a lesser degree. This may help explain why none of the clones could be reliably identified in the normal endometrium of Patient 43.

The figure below shows that, like for patients 39 and 42, multiple instances of hotspot mutations in *PIK3CA* and *KRAS* could be identified. However, no more than two endometriosis barcode mutations could be called in any individual segment. Clone #2 in the figure below thus exemplifies a negative finding.

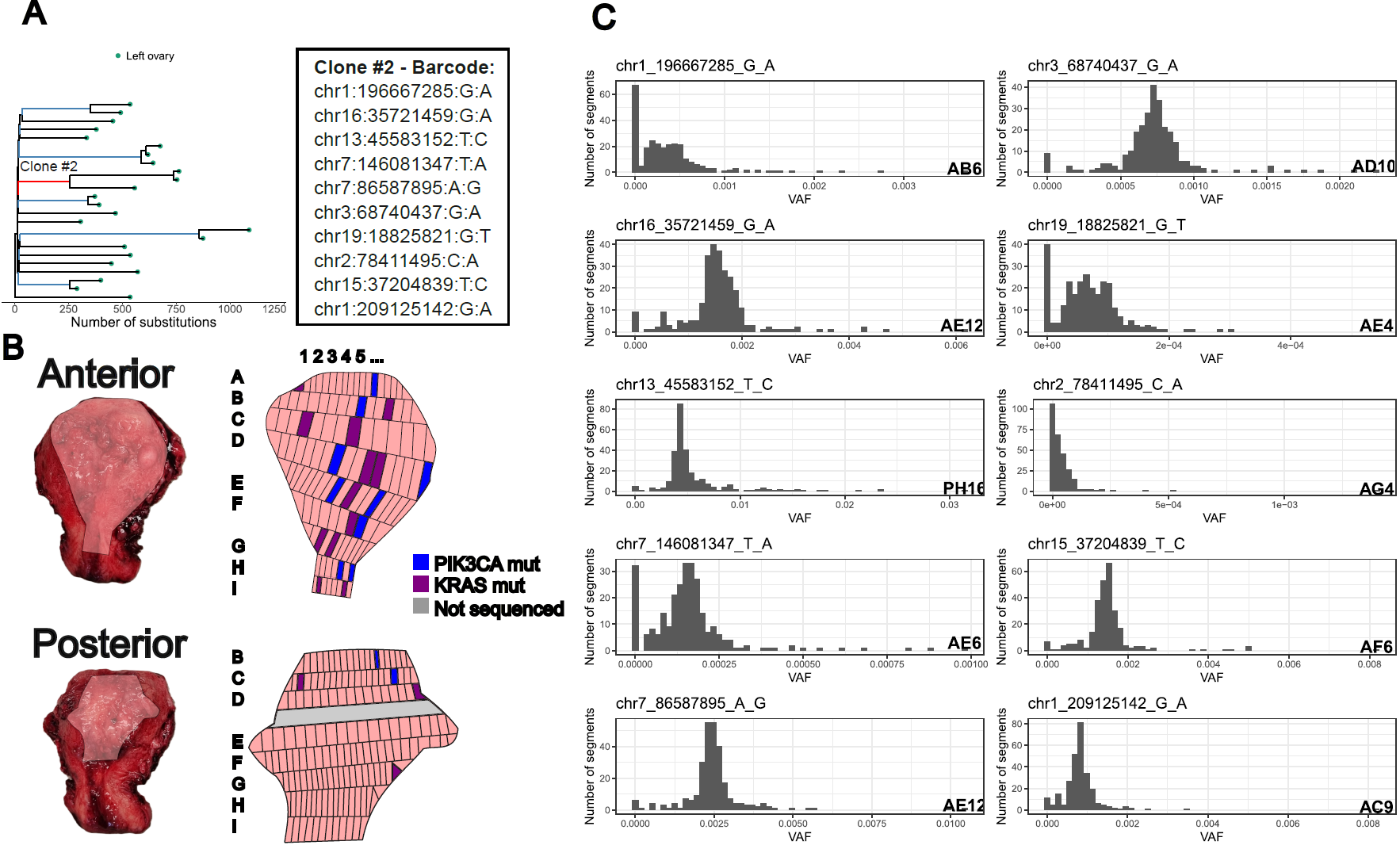

**Screening for endometriosis clones in the normal endometrium of patient 43.** Six endometriosis clones were selected for the screen, highlighted in blue and red in panel A. Results are shown for Clone #2. A) a 10 mutation barcode was selected from the most ancestral branch possible. B) The whole endometrium was isolated and divided into 274 small (approximately 2x2mm) segments. The gray line of the posterior half was removed for clinical work and was unavailable for the study. C) Histograms showing the variant allele fraction (VAF) of each of the ten mutations comprising the barcode for Clone #2 and highlighting the endometrial segment with the highest VAF. While some segments are outliers on the VAF distributions, these are often segments with low coverage at the site and/or segments where all the reads reporting the mutation map to one strand. Mutations passing filters are sporadically called but no segment shows multiple mutations from the same clone barcode.

Supplementary Figures

Fig. S1. Coverage and variant-allele fractions for microdissections stratified by endometriosis lesional status. Does not include samples from the screening of whole uteri.

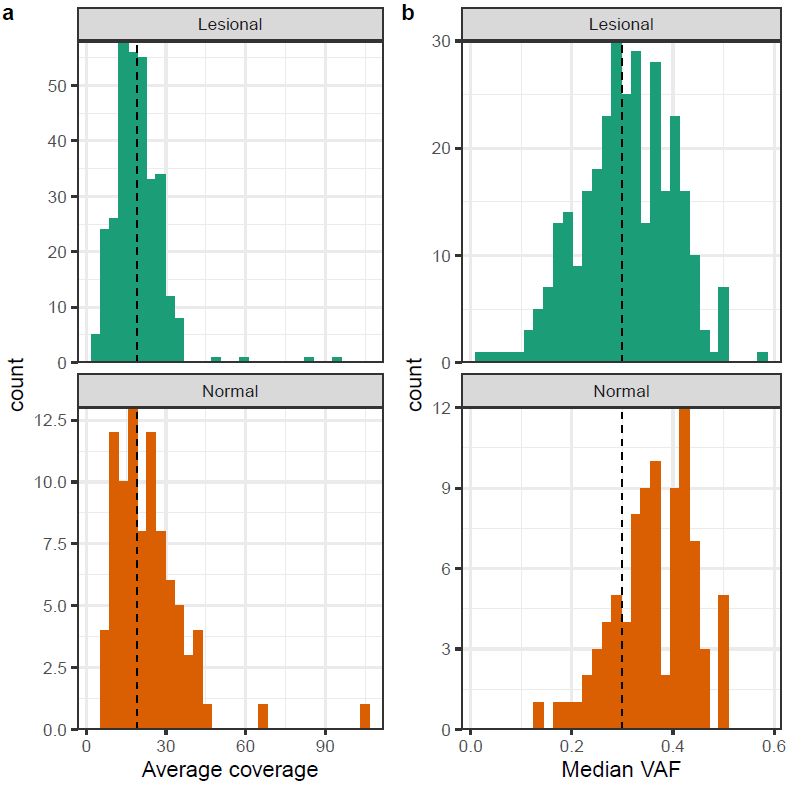
a) A histogram showing the sequencing coverage of all microdissections stratified into lesional dissections and normal glands. The dashed line shows the median sequencing coverage for the cohort as a whole (18.9X). b) A histogram showing the median variant allele fraction (VAF) for all micro dissected samples. The dashed line shows VAF=0.3 for ease of reading.

Fig. S2. Sensitivity estimates of the mutation calling pipeline at different coverages.

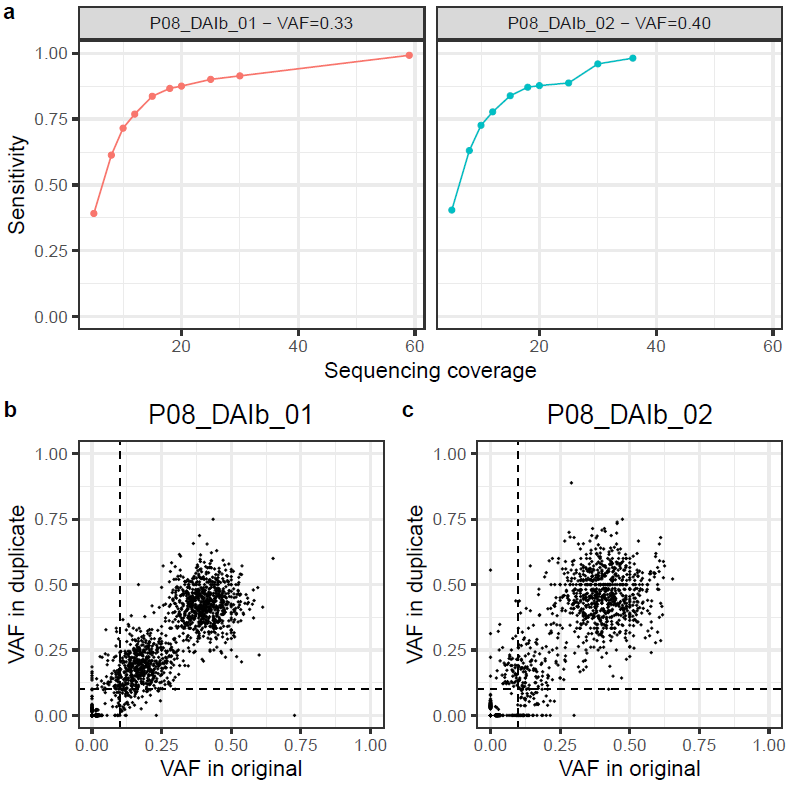

Sequencing of near-duplicate samples. We dissected adjacent histological sections of the same two endometrial glands (glandIDs: P08_DAIb_01 and P08_DAIb_02) into two wells. We created independent sequencing libraries for each replicate sample. a) Sensitivity of the mutation calling pipeline as a function of coverage in down-sampled samples. b and c) the variant allele fraction of mutations detected using the maximum possible coverage.

Fig. S3. Mutational signatures extracted by HDP.

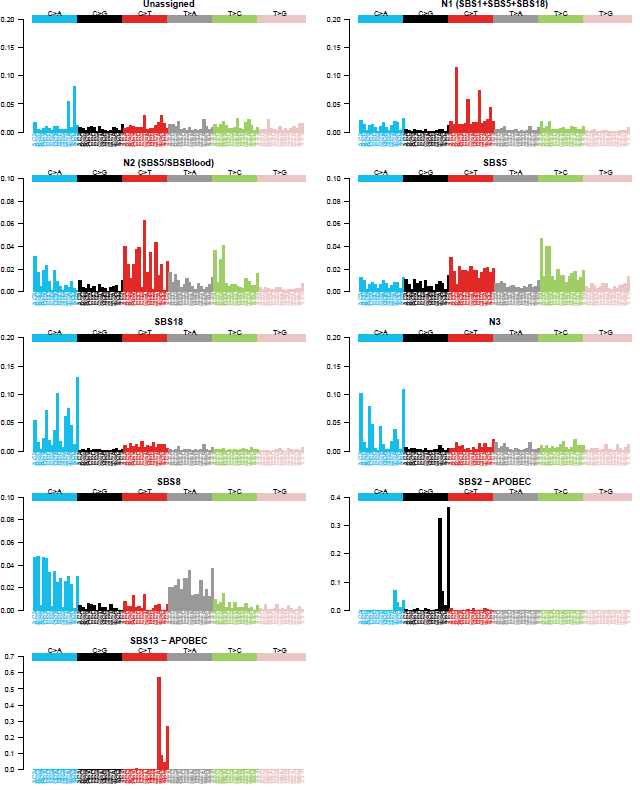

Mutational signatures identified by hierarchical Dirichlet process. Component N1 was divided into SBS1, SBS5 and SBS18 using an expectation maximization algorithm. Component N2 was merged with SBS5. Component N3 was merged with the “Unassigned” component

Fig. S4. Phylogenetic tree for Patient 1.

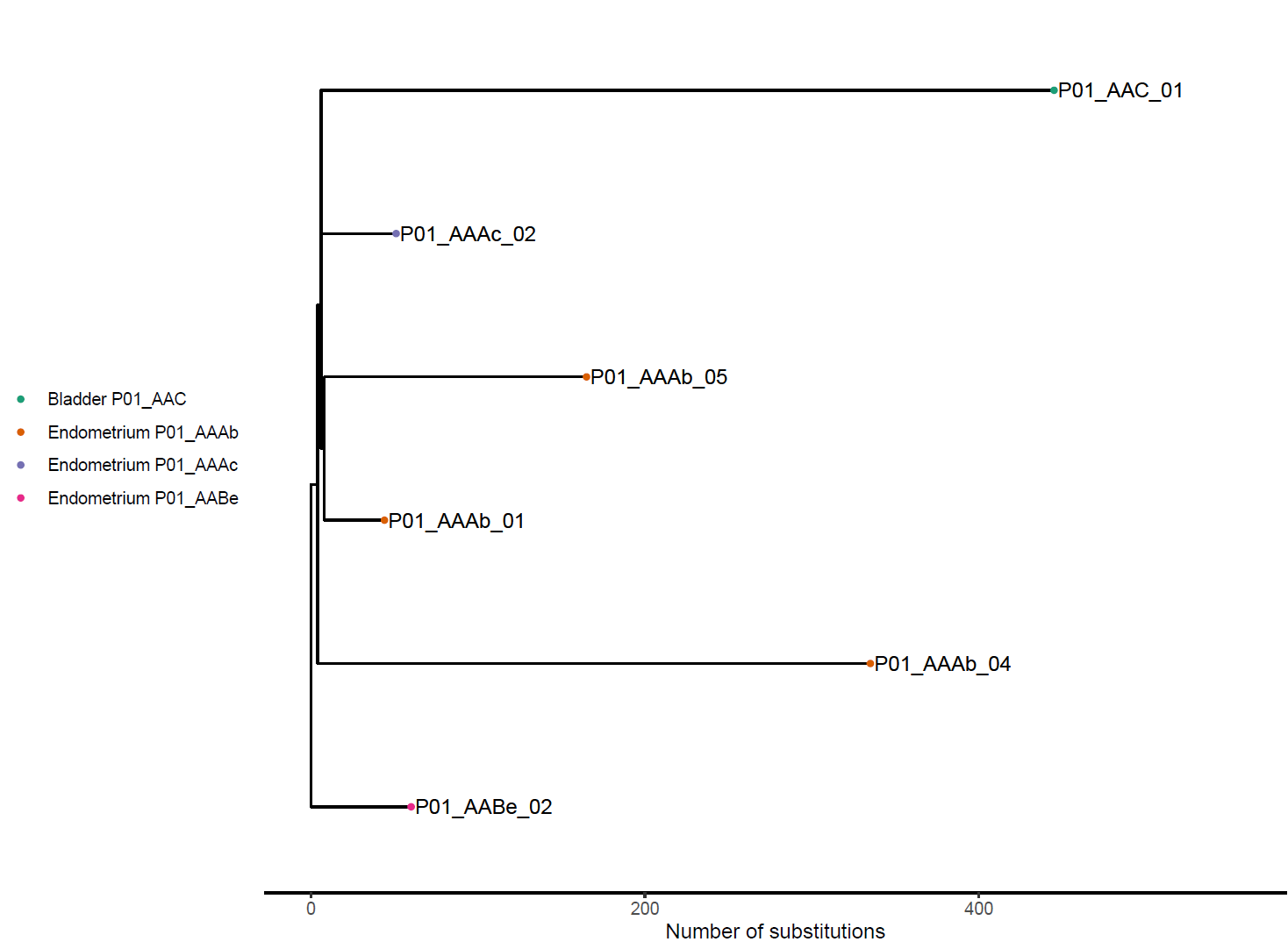

Fig. S5. Phylogenetic tree for Patient 2.

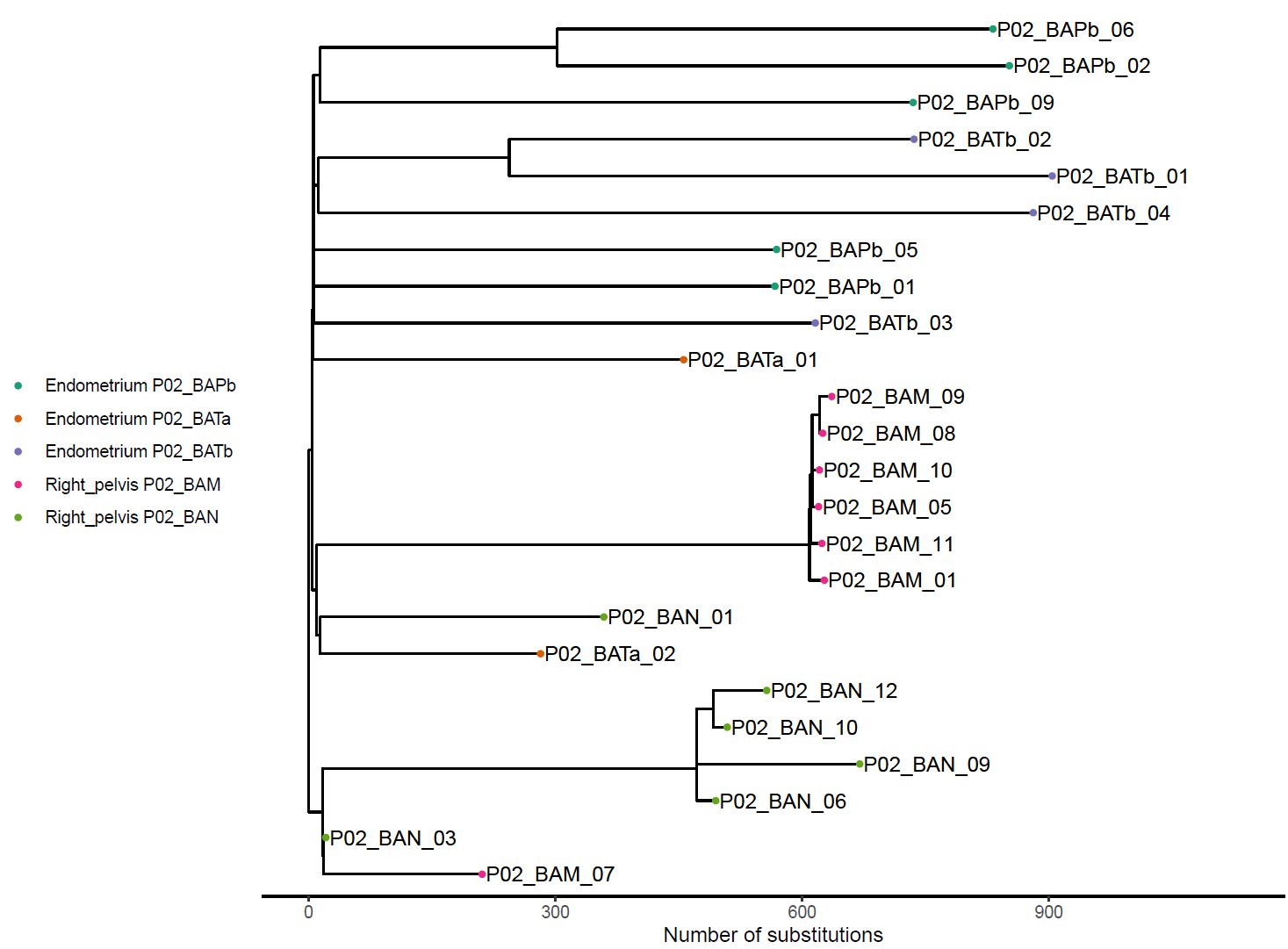

Fig. S6. Phylogenetic tree for Patient 3.

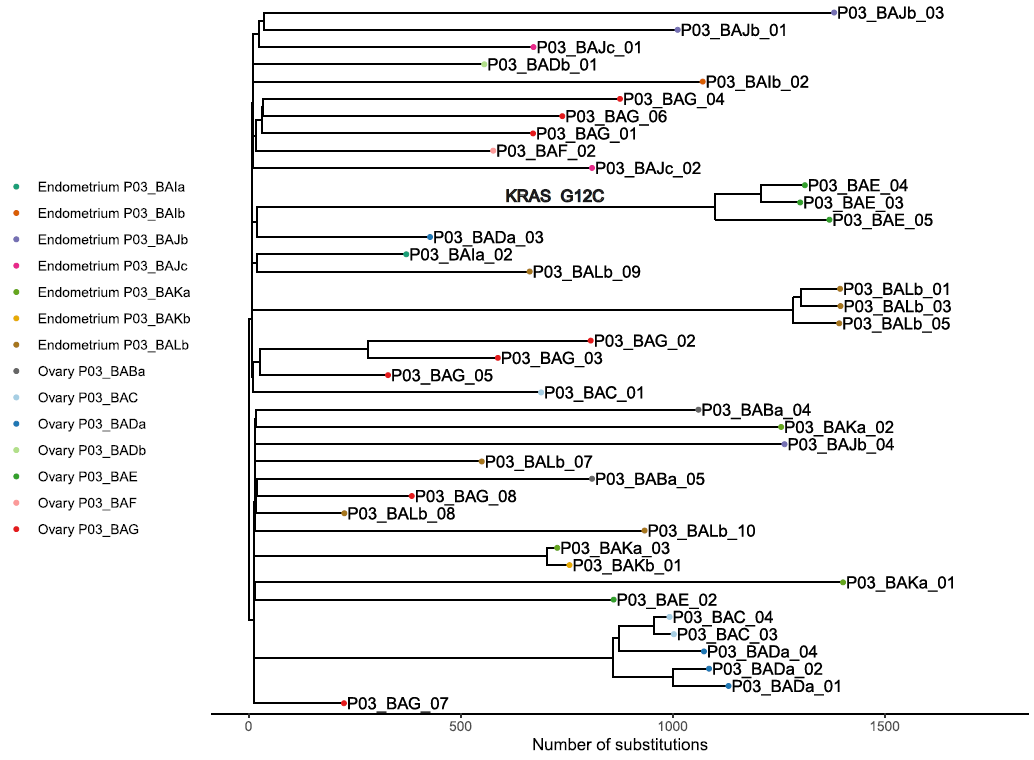

Fig. S7 Phylogenetic tree for Patient 5.

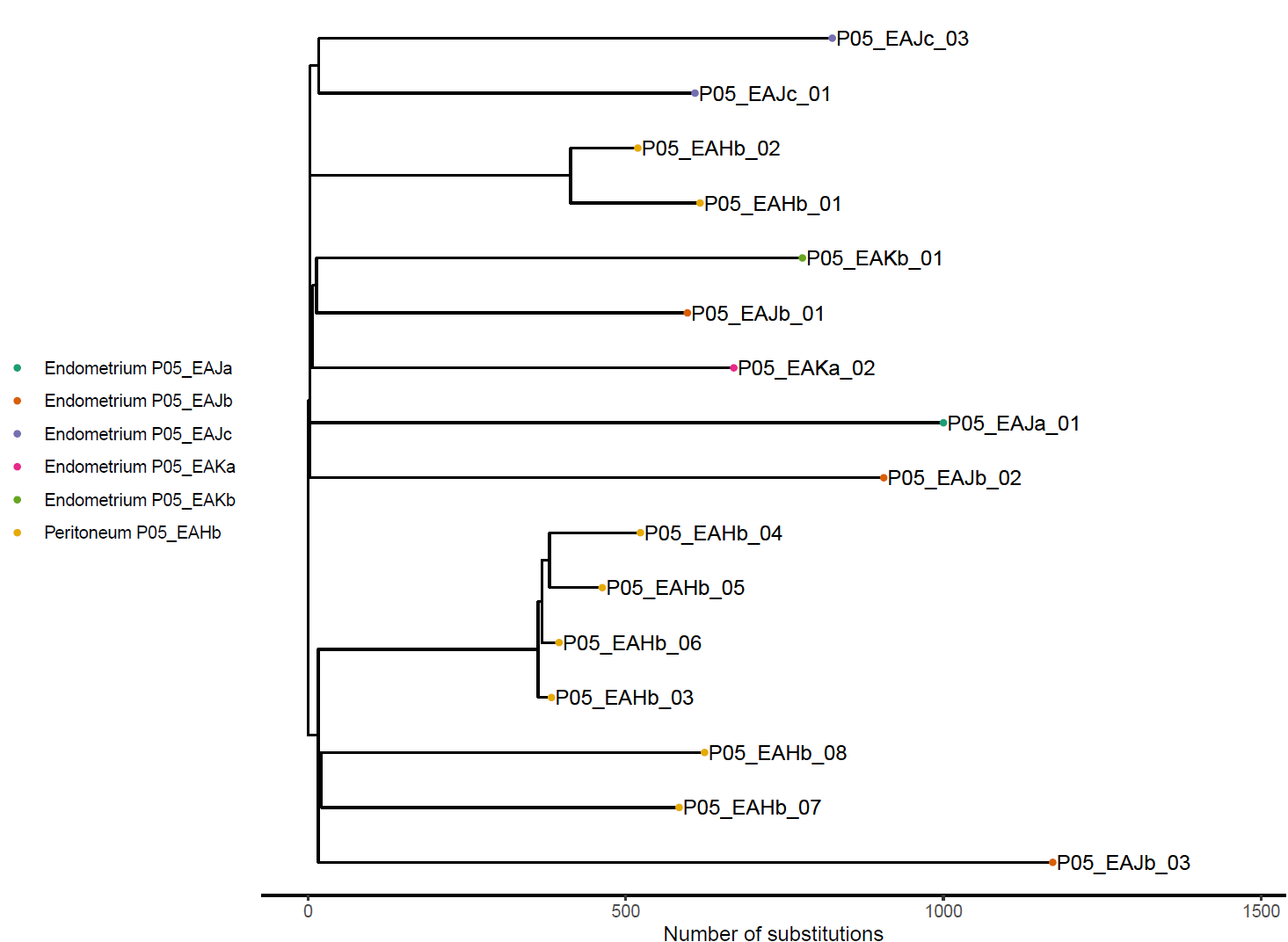

Fig. S7. Phylogenetic tree for Patient 7.

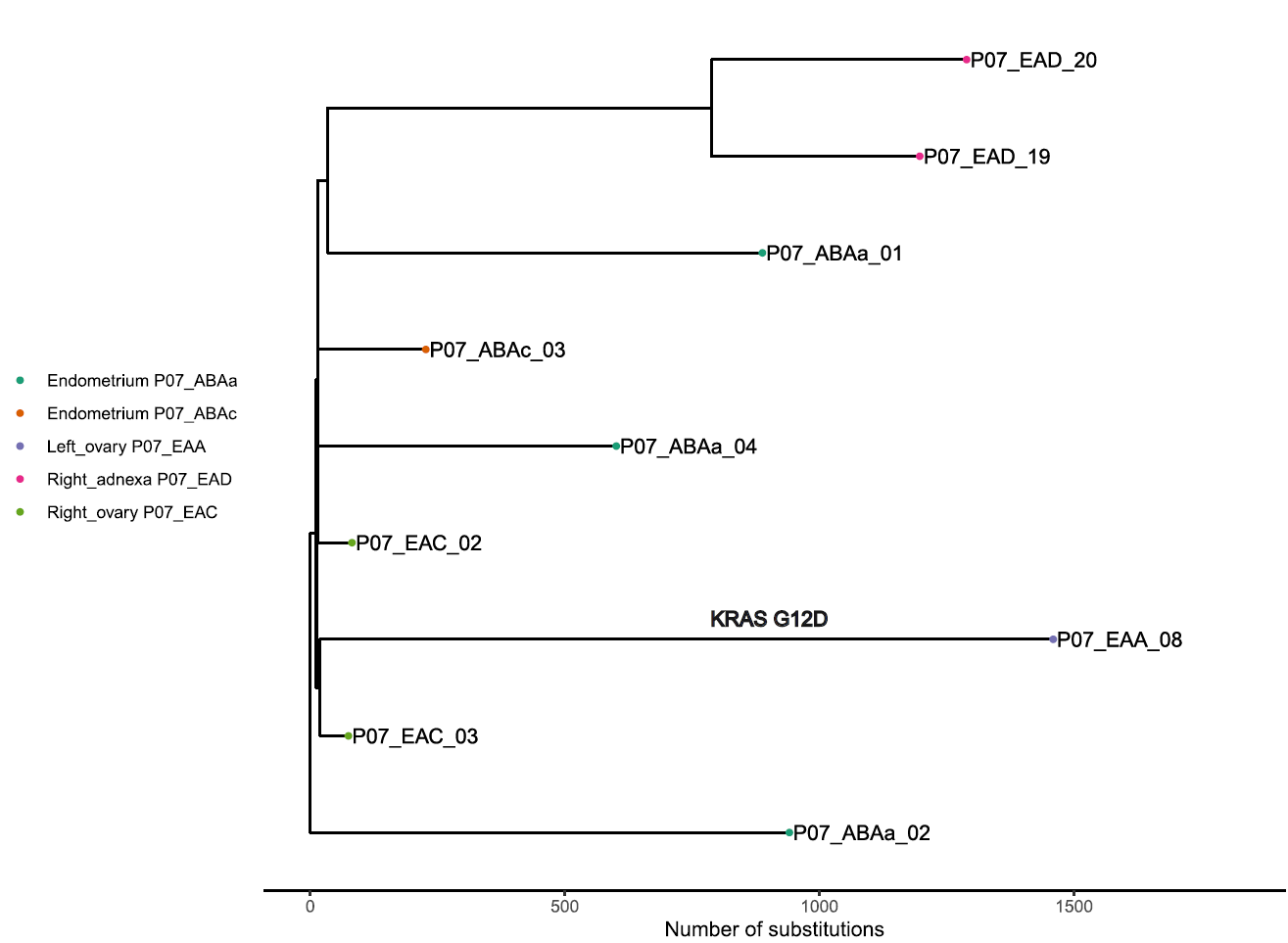

Fig. S8. Phylogenetic tree for Patient 8.

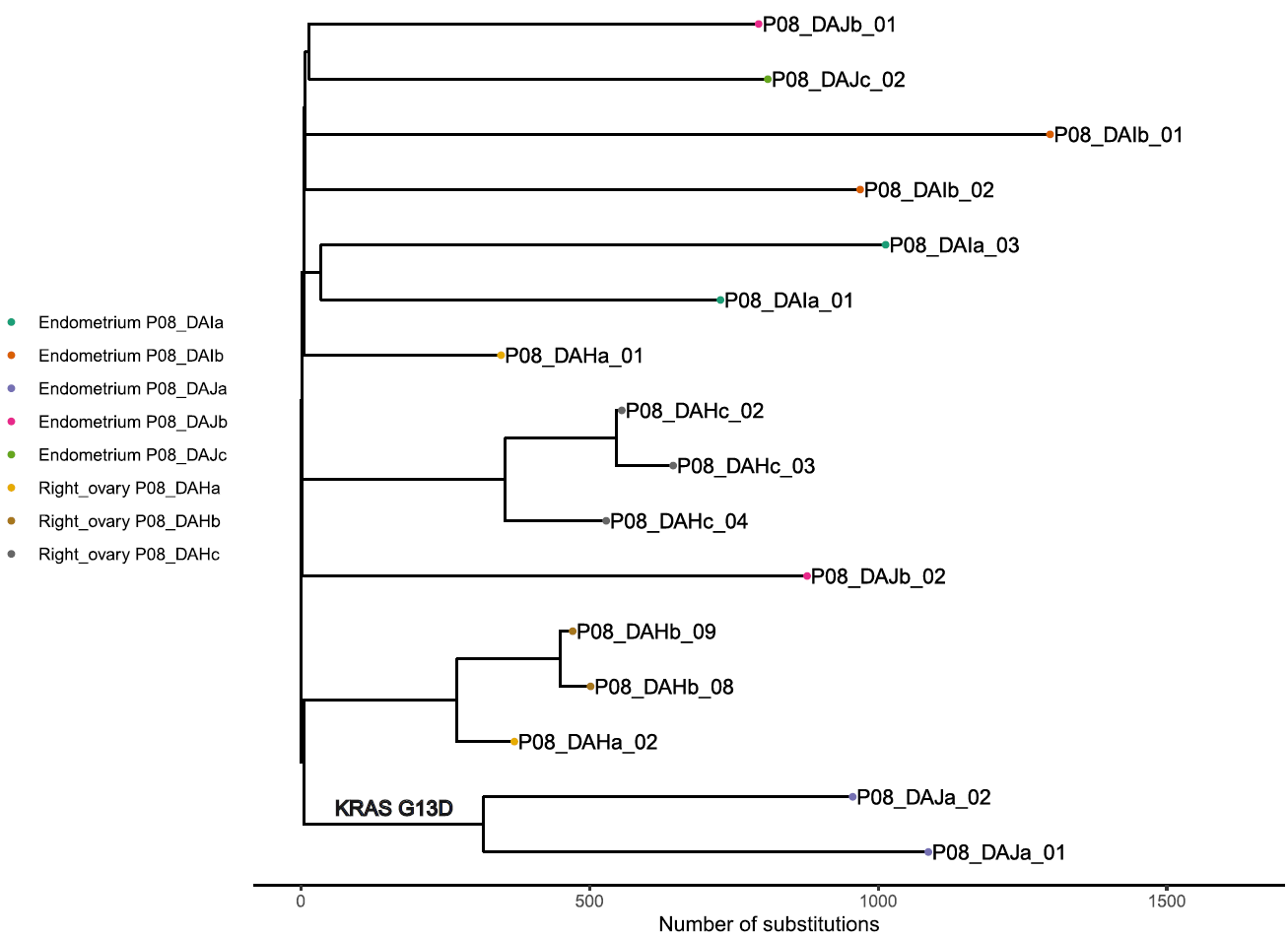

Fig. S9. Phylogenetic tree for Patient 9.

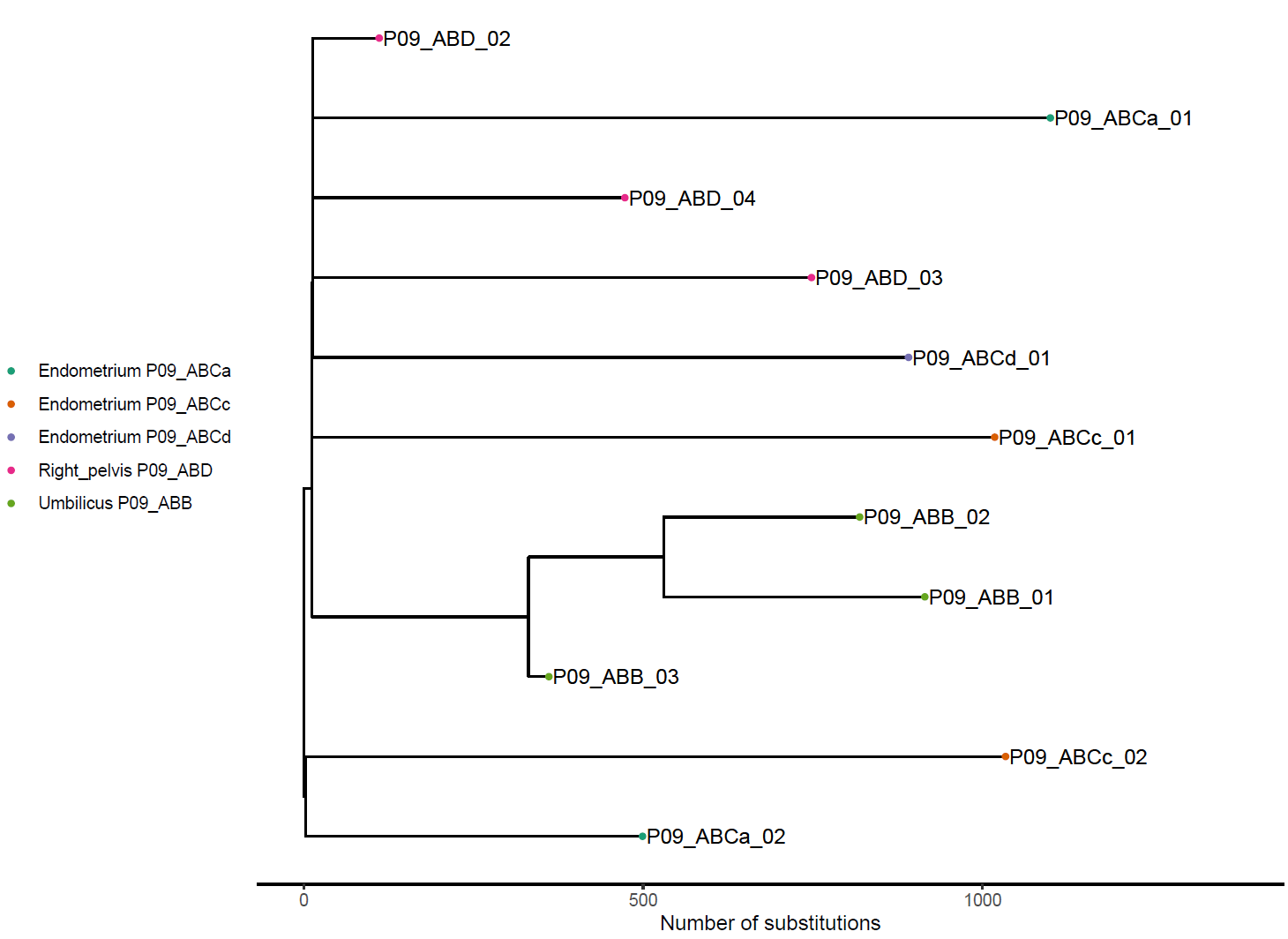

Fig. S10. Phylogenetic tree for Patient 10.

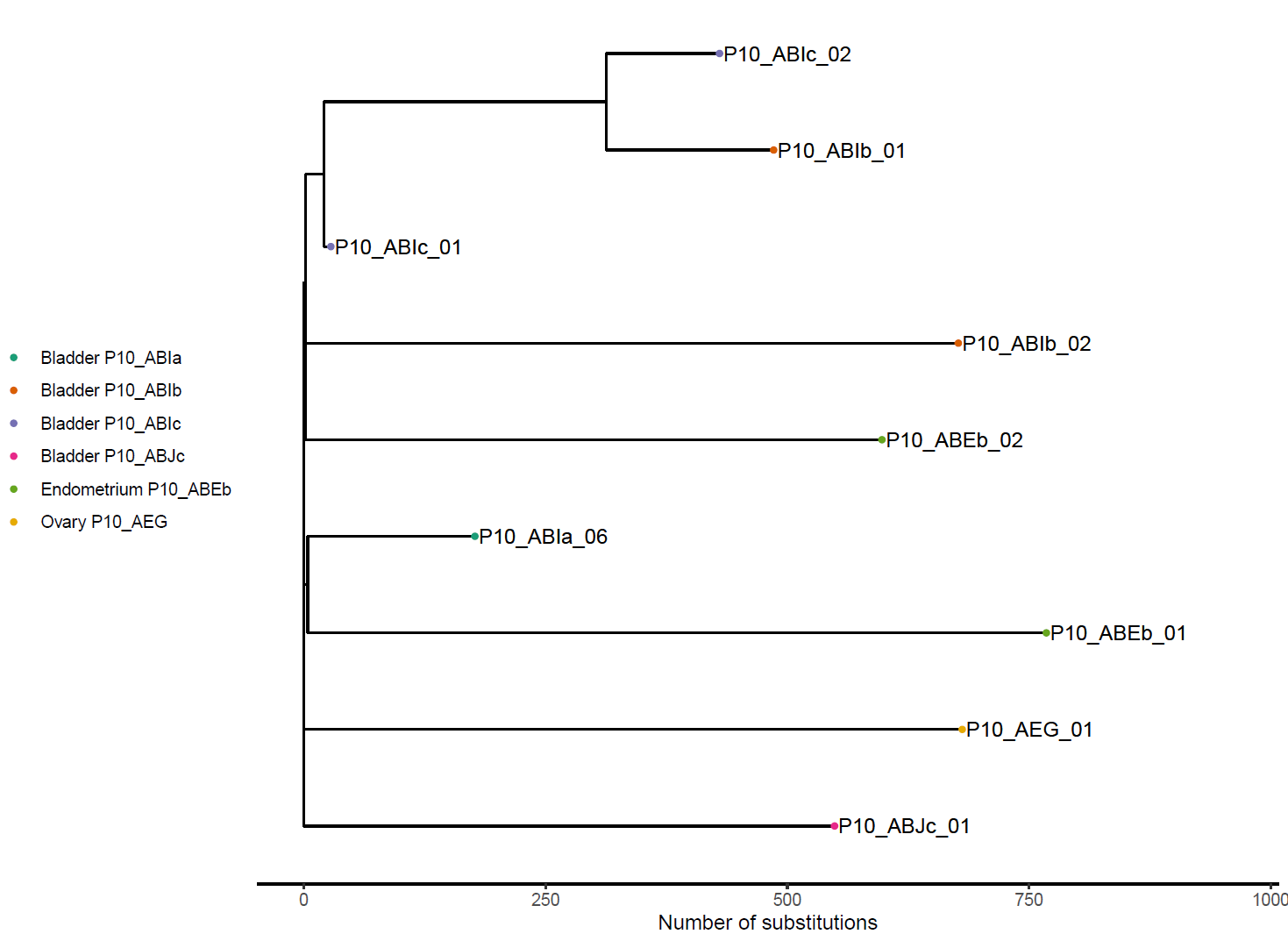

Fig. S11. Phylogenetic tree for Patient 15.

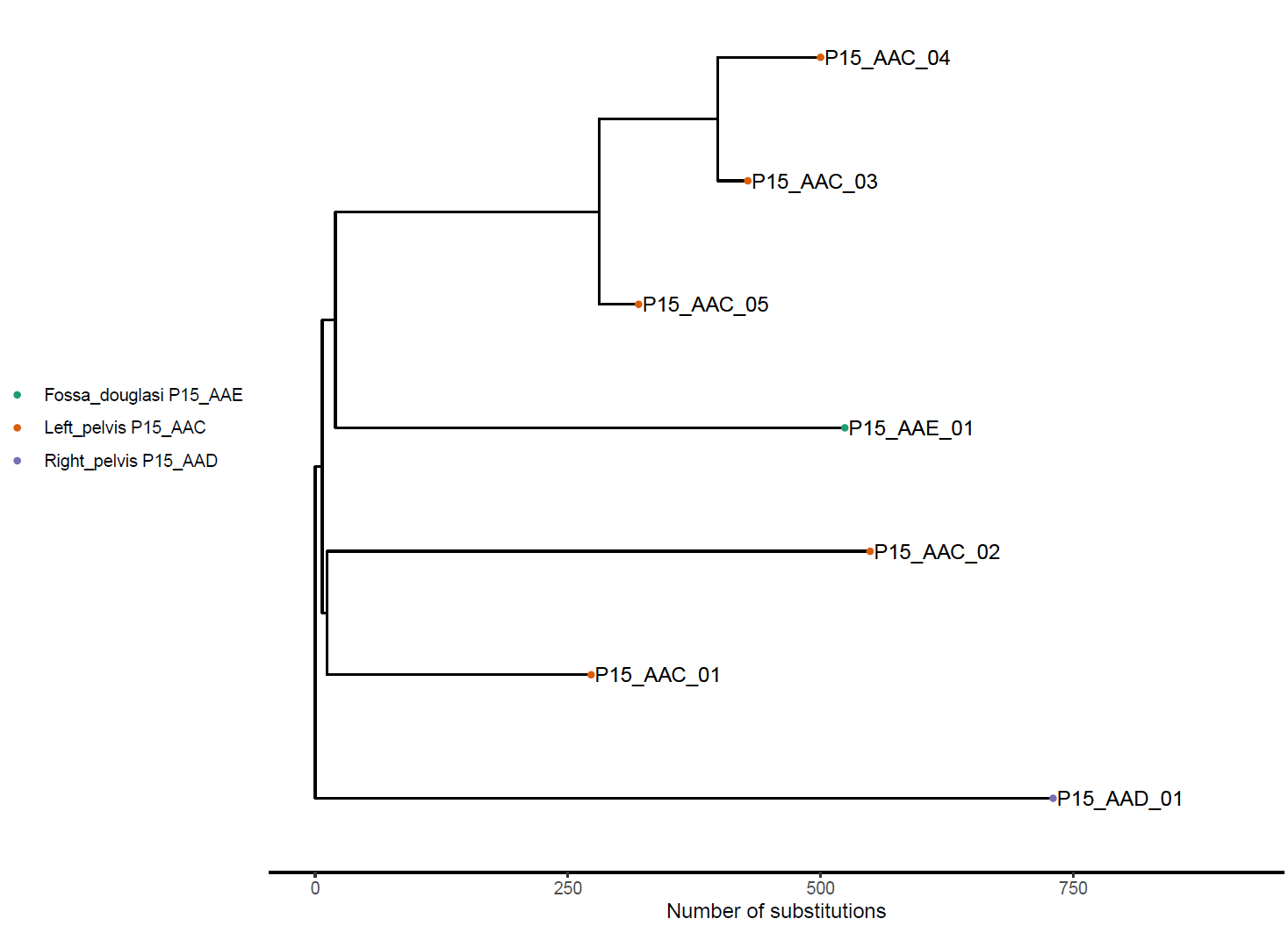

Fig. S12. Phylogenetic tree for Patient 17.

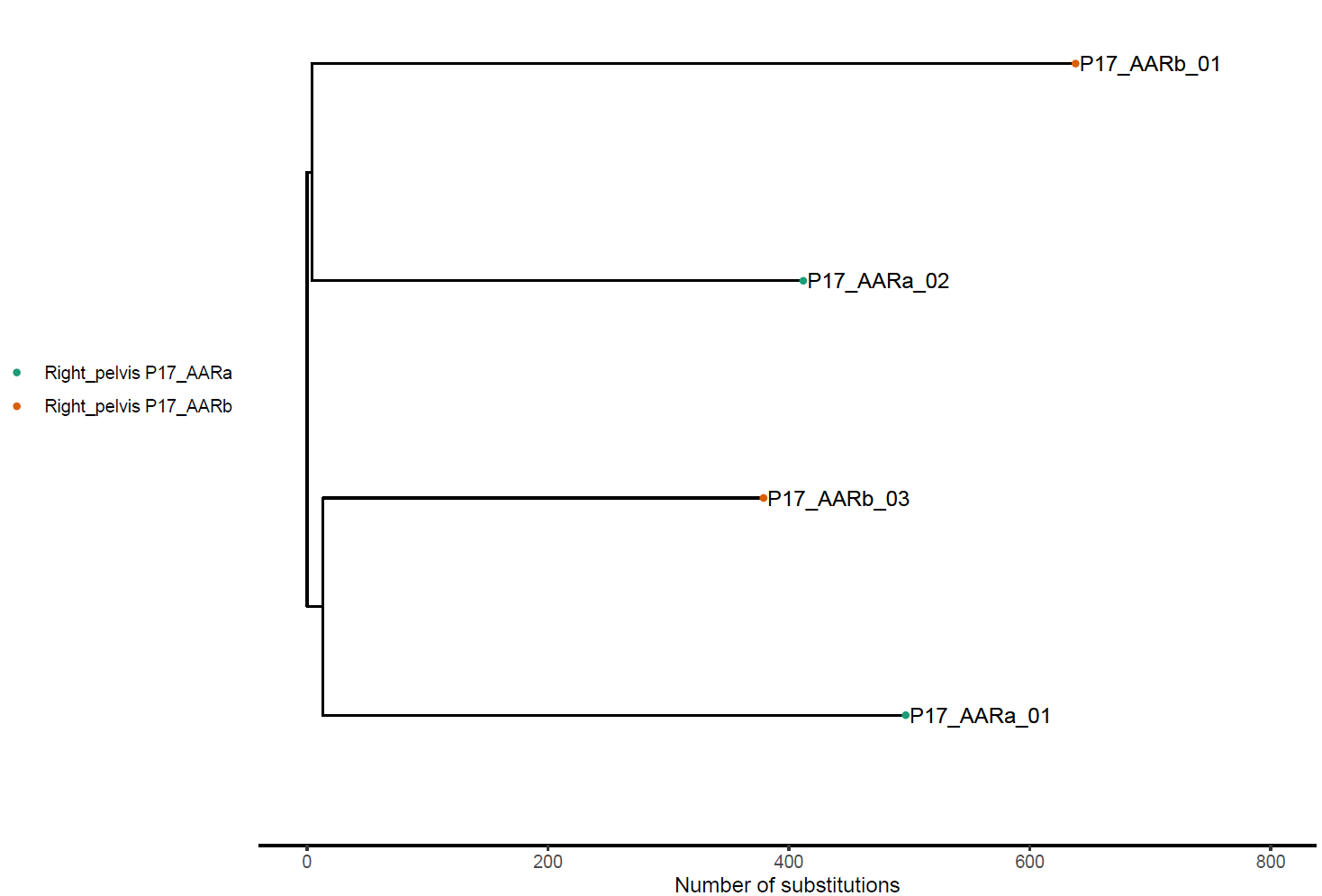

Fig. S13. Phylogenetic tree for Patient 18.

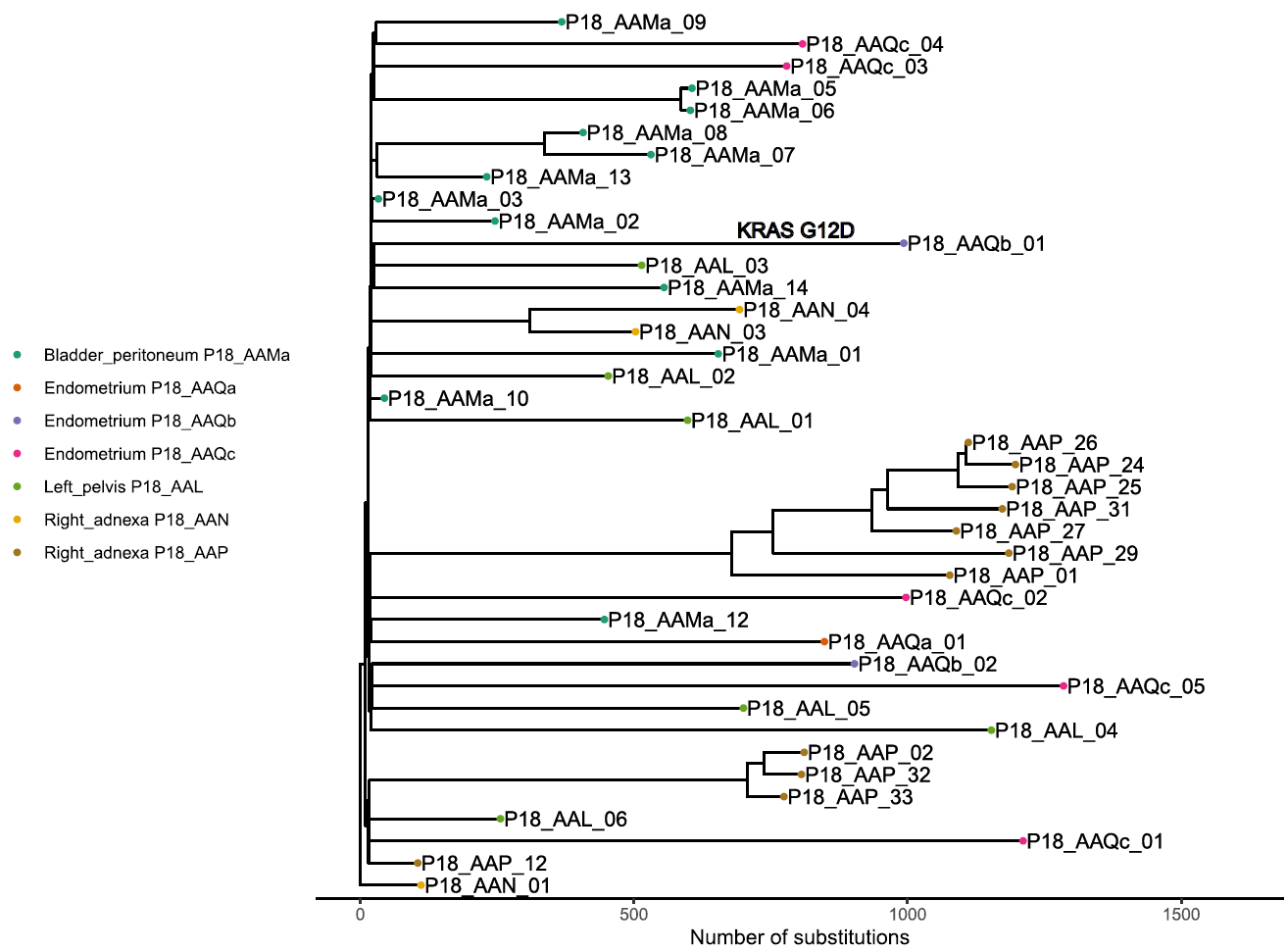

Fig. S14. Phylogenetic tree for Patient 19.

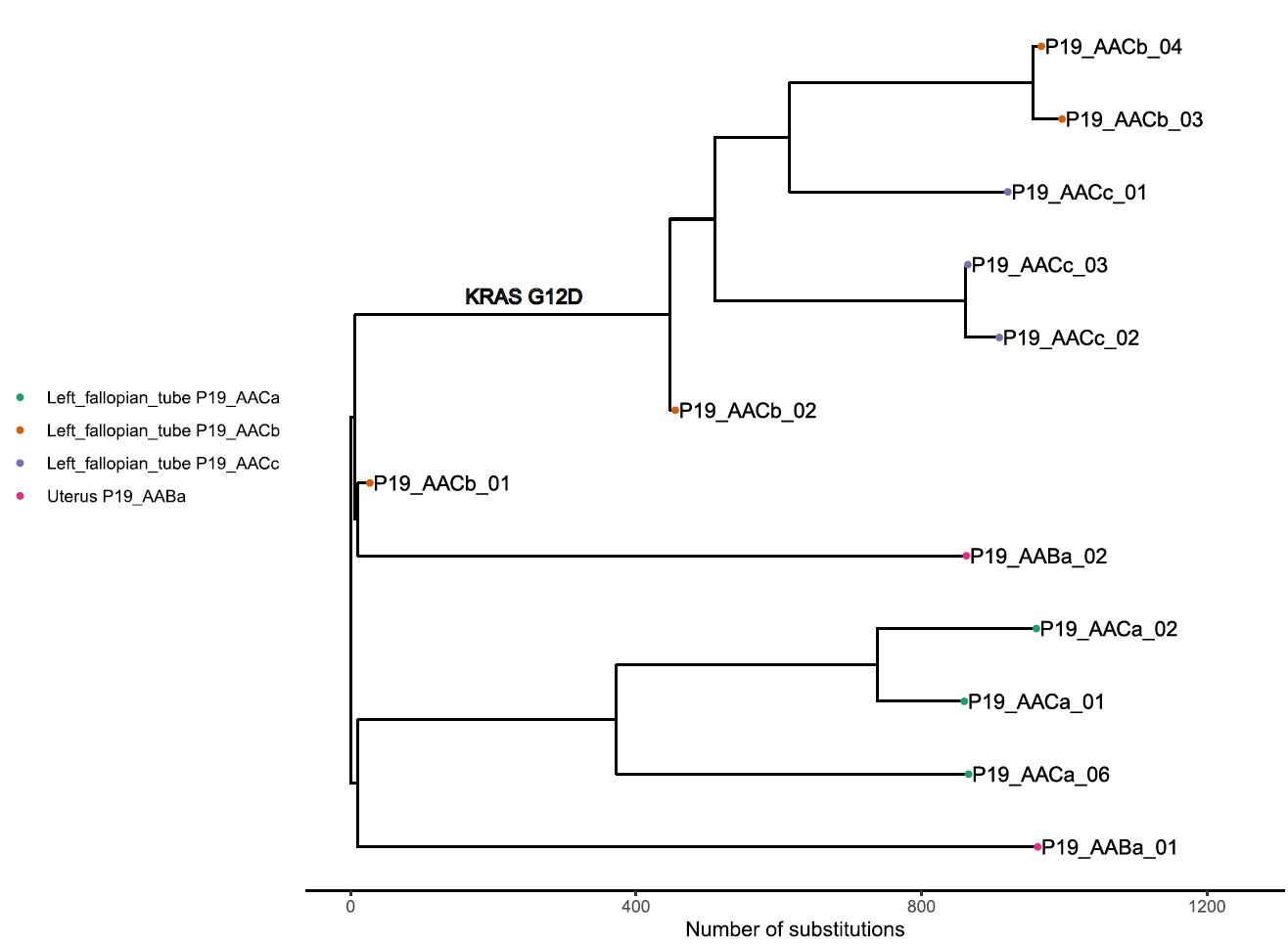

Fig. S15. Phylogenetic tree for Patient 21.

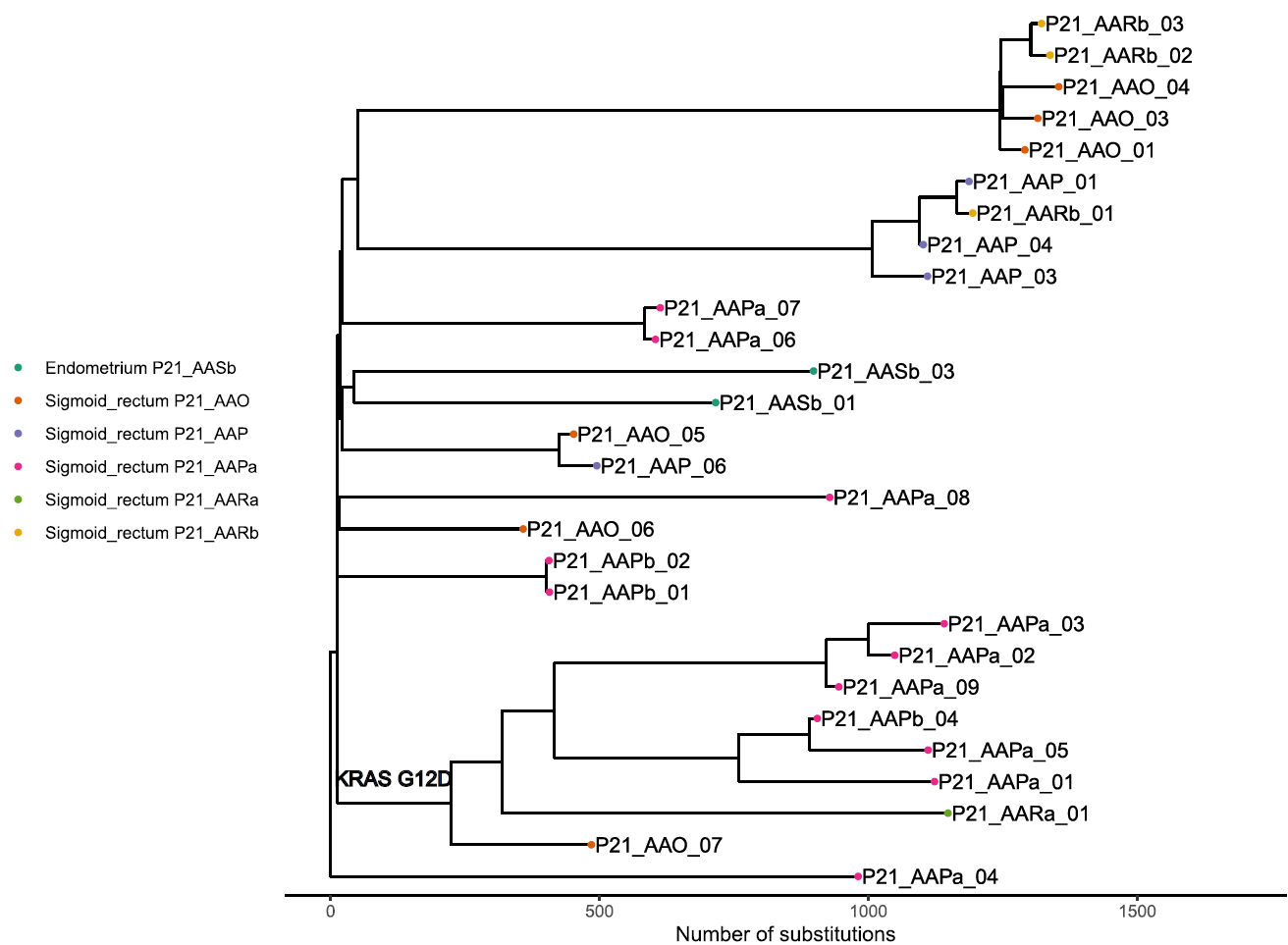

Fig. S16. Phylogenetic tree for Patient 23.

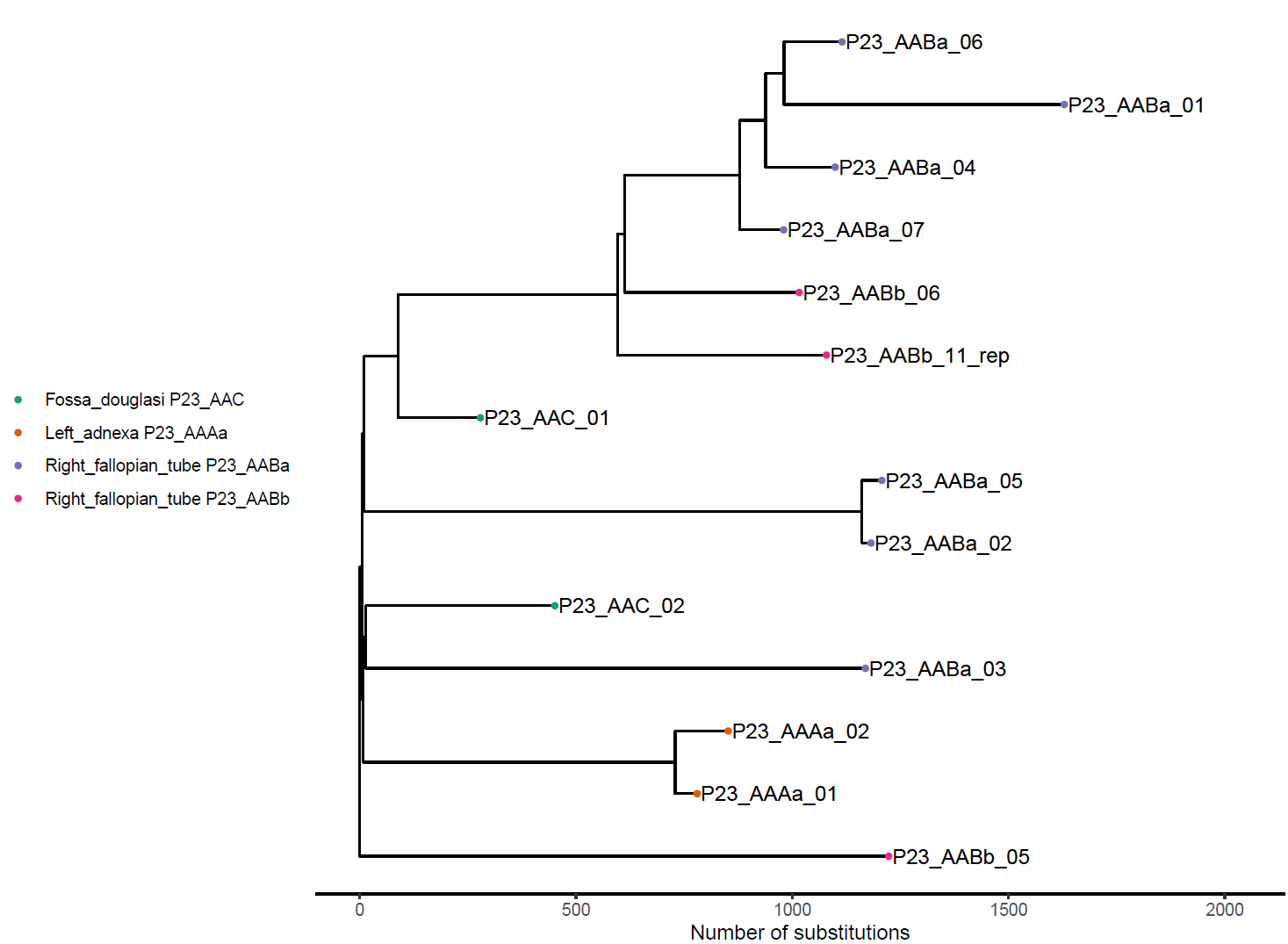

Fig. S17. Phylogenetic tree for Patient 25.

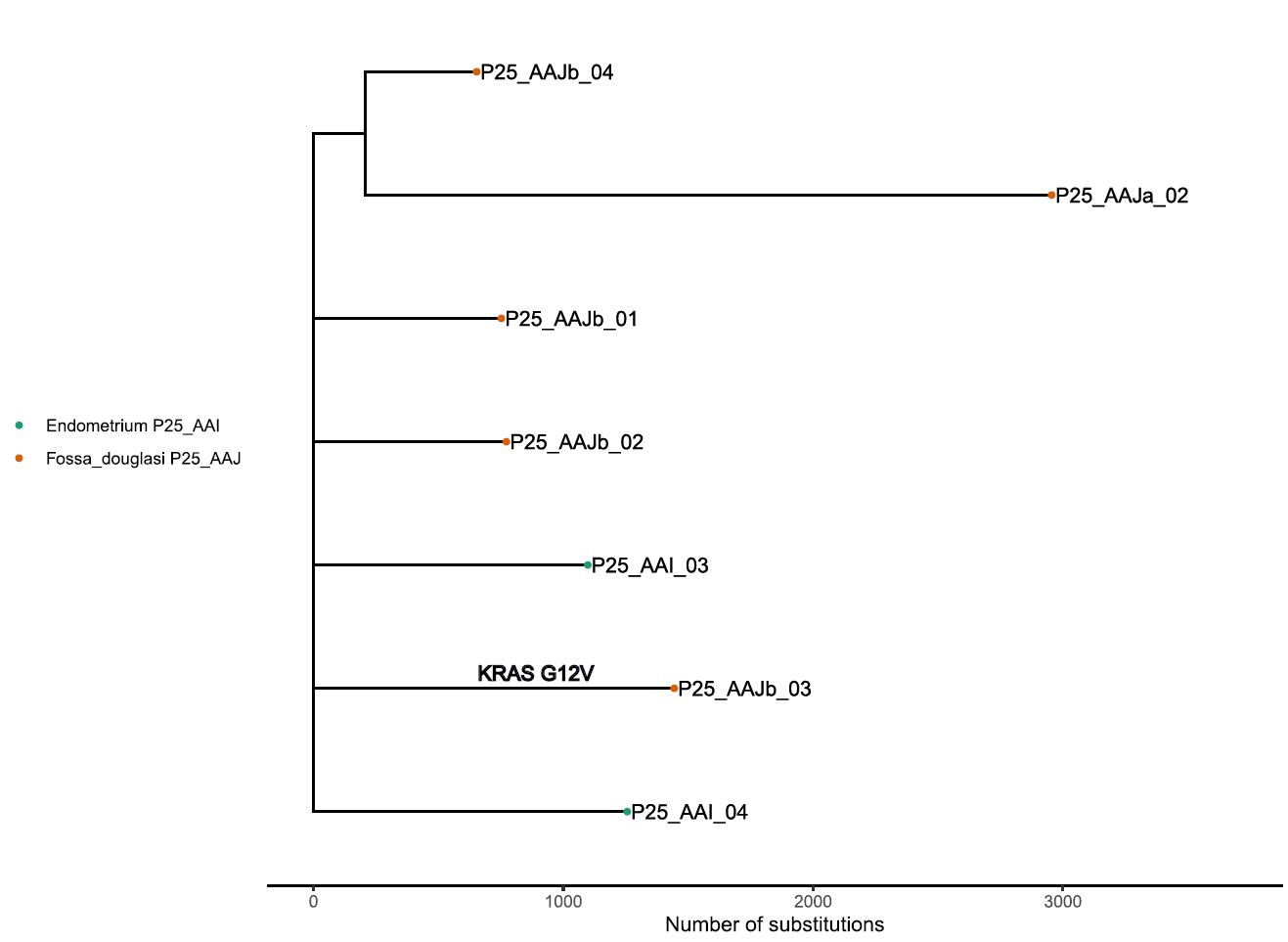

Fig. S18. Phylogenetic tree for Patient 28.

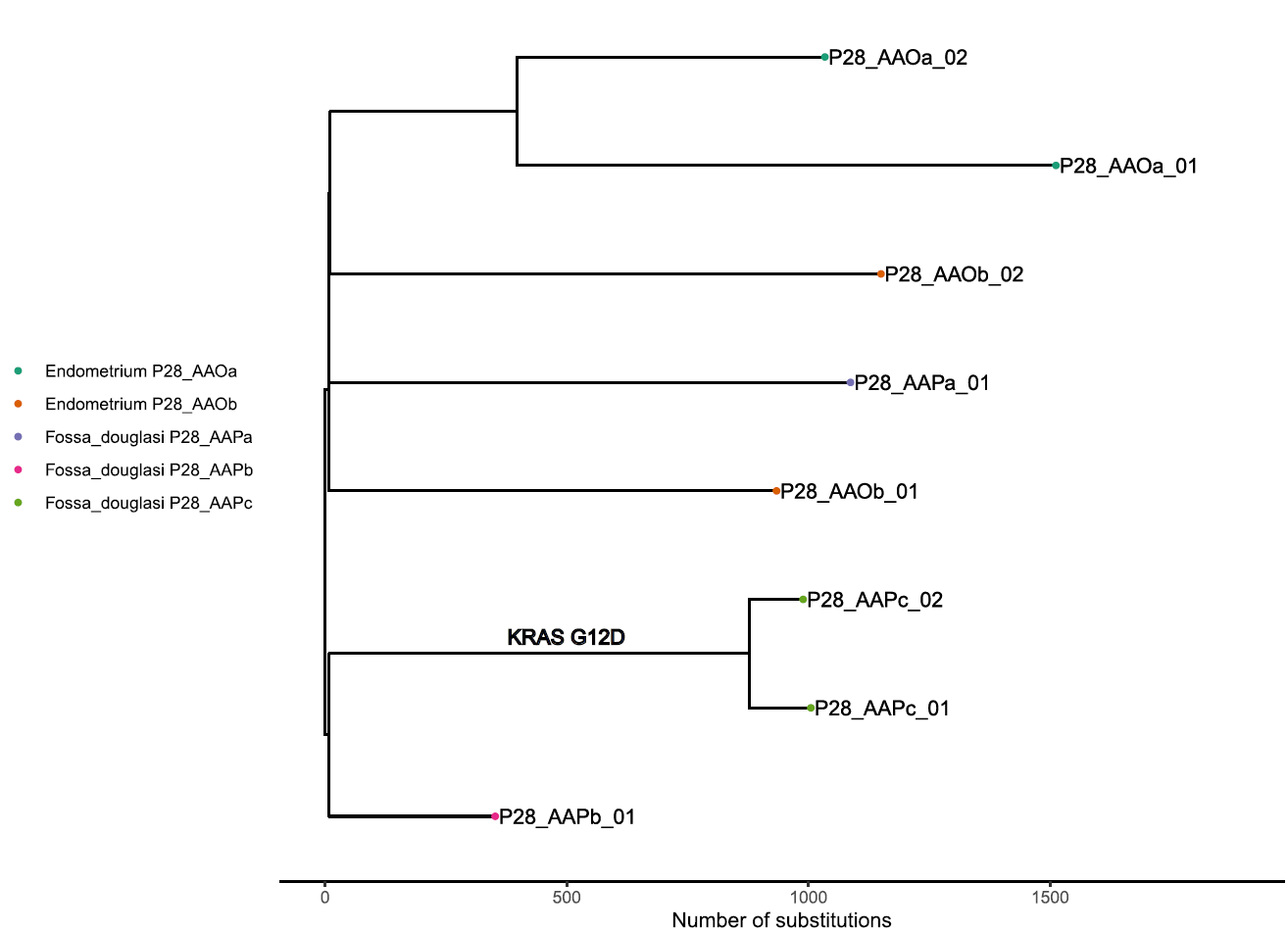

Fig. S19. Phylogenetic tree for Patient 30.

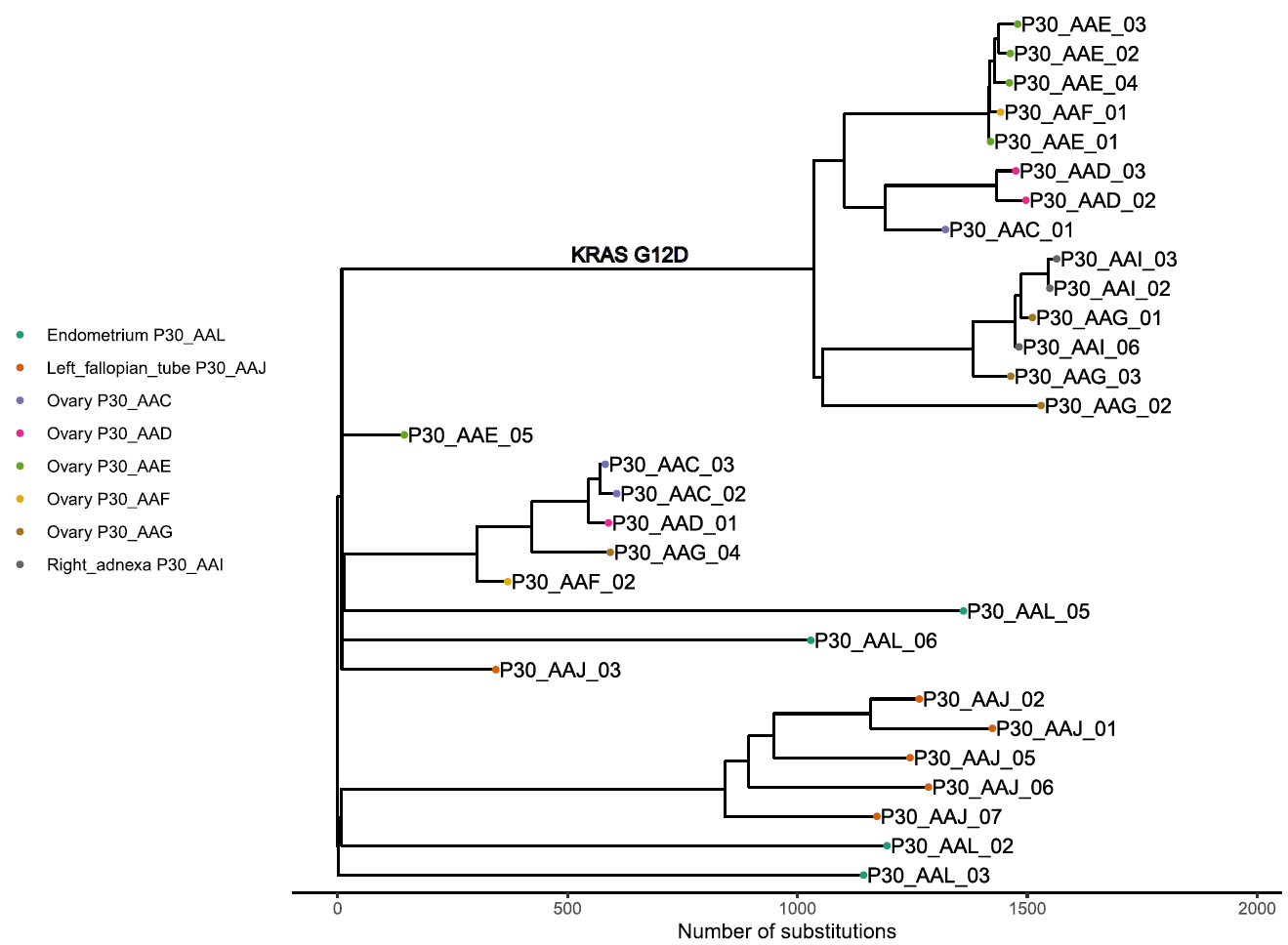

Fig. S20. Phylogenetic tree for Patient 32.

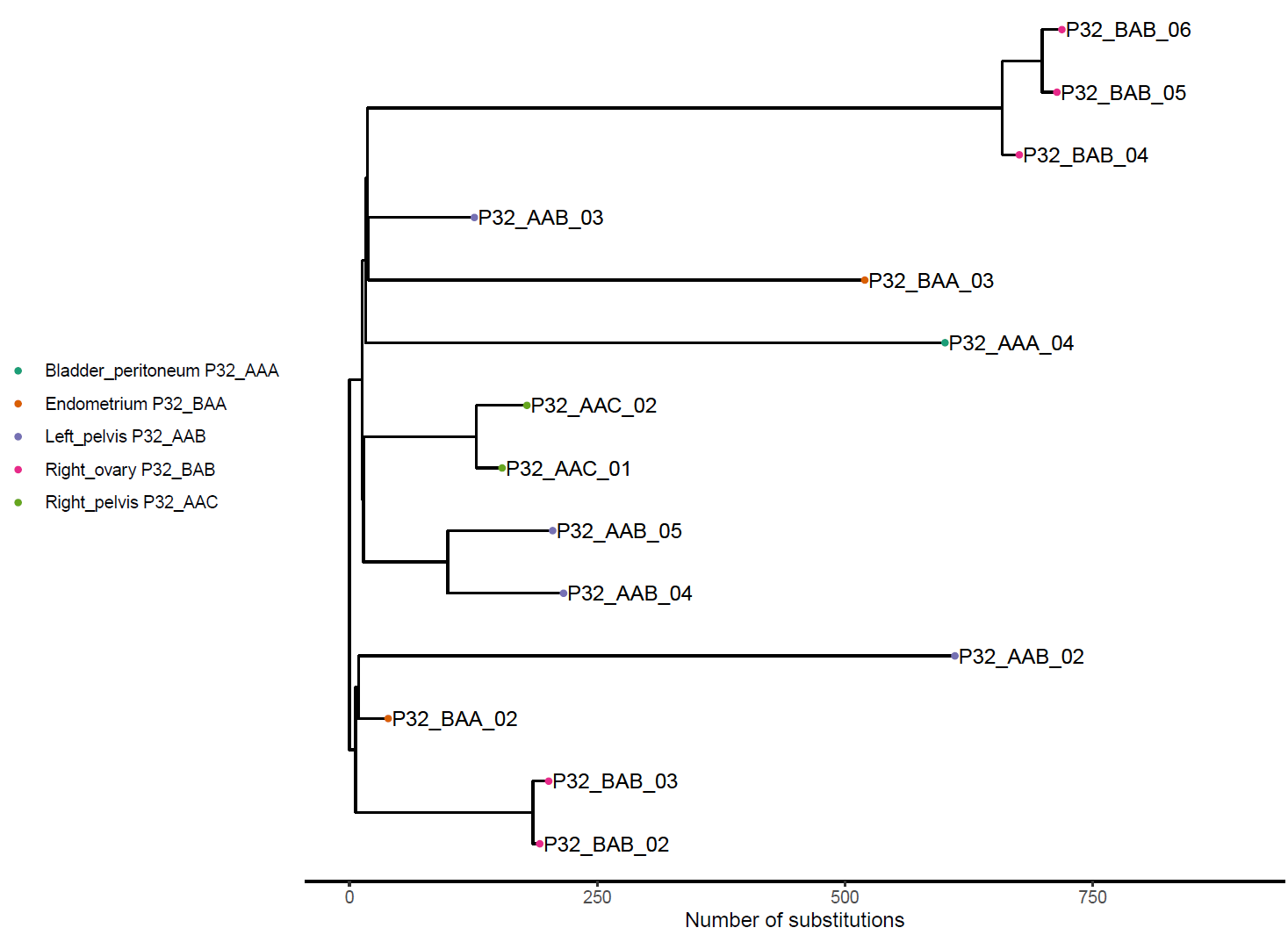

Fig. S21. Phylogenetic tree for Patient 35.

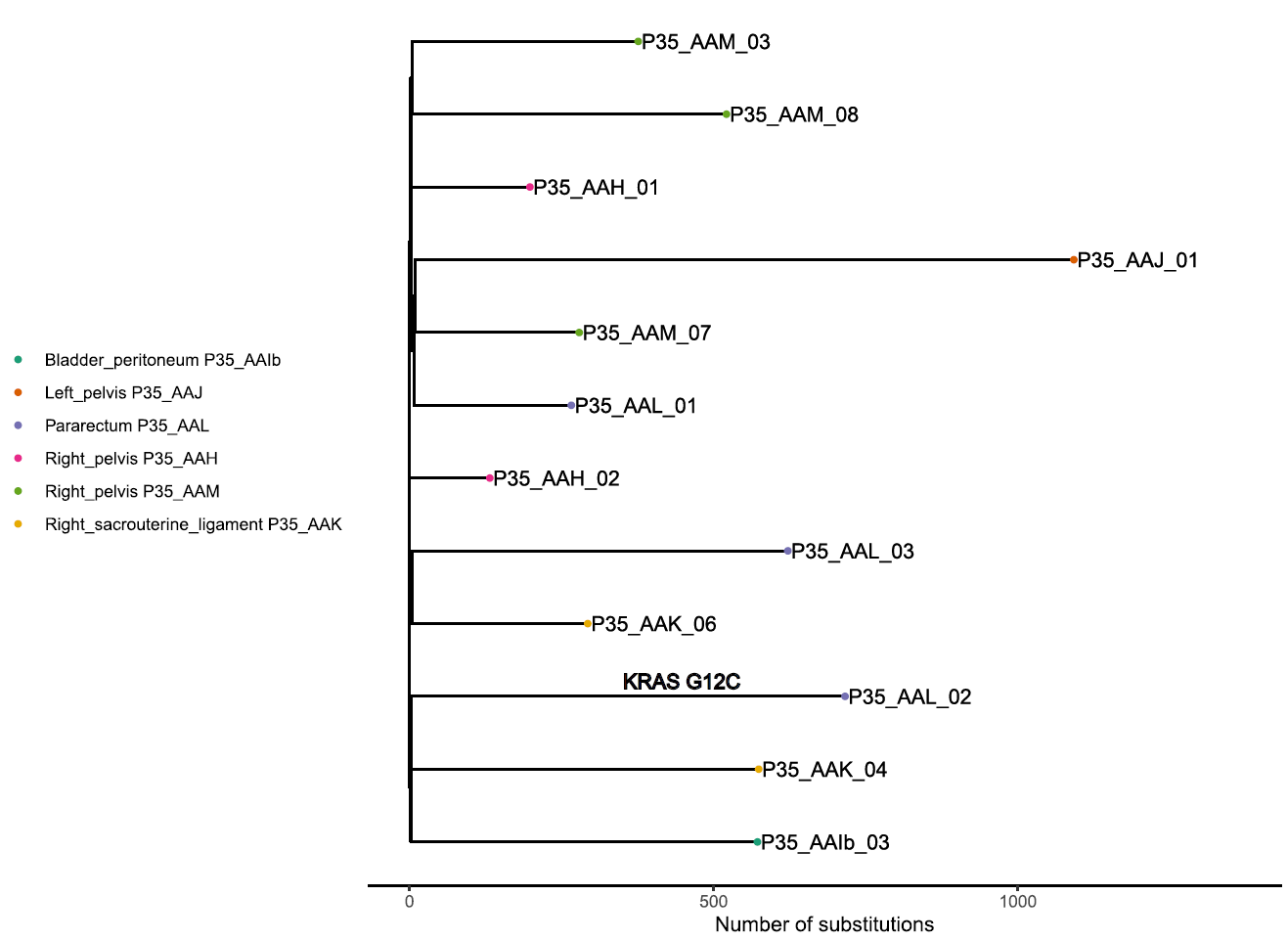

Fig. S22. Phylogenetic tree for Patient 37.

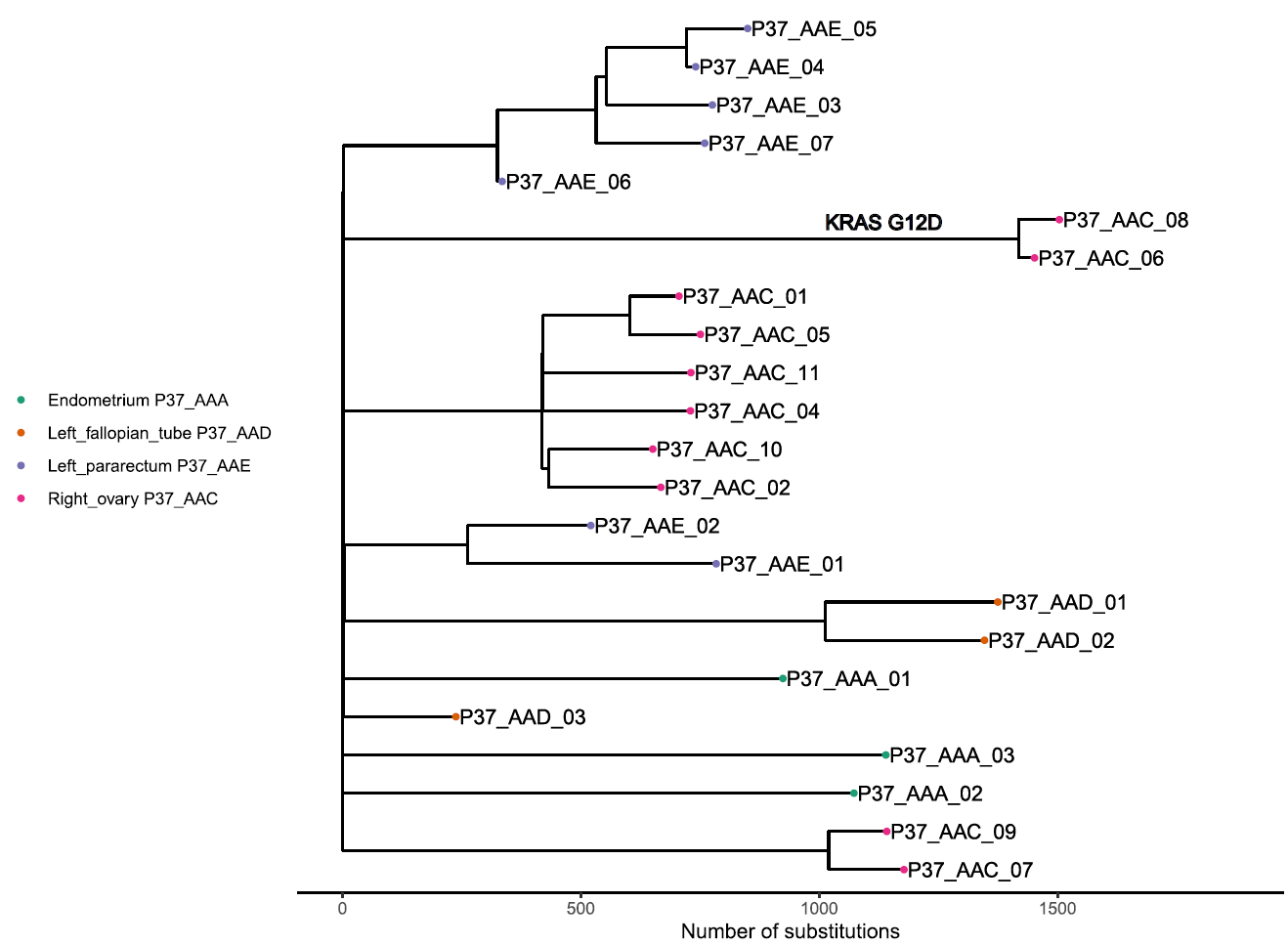

Fig. S23. Phylogenetic tree for Patient 39.

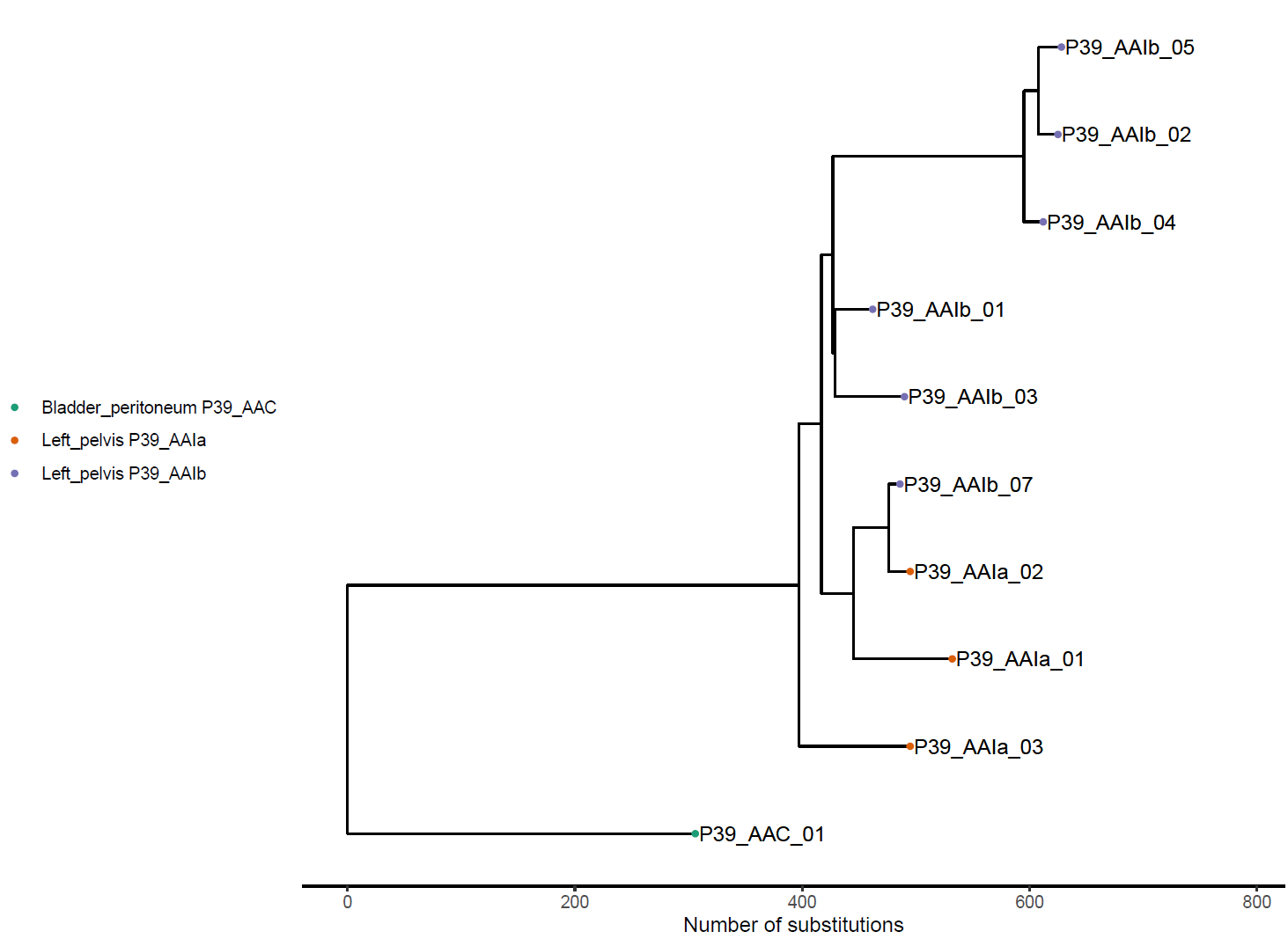

Fig. S24. Phylogenetic tree for Patient 40.

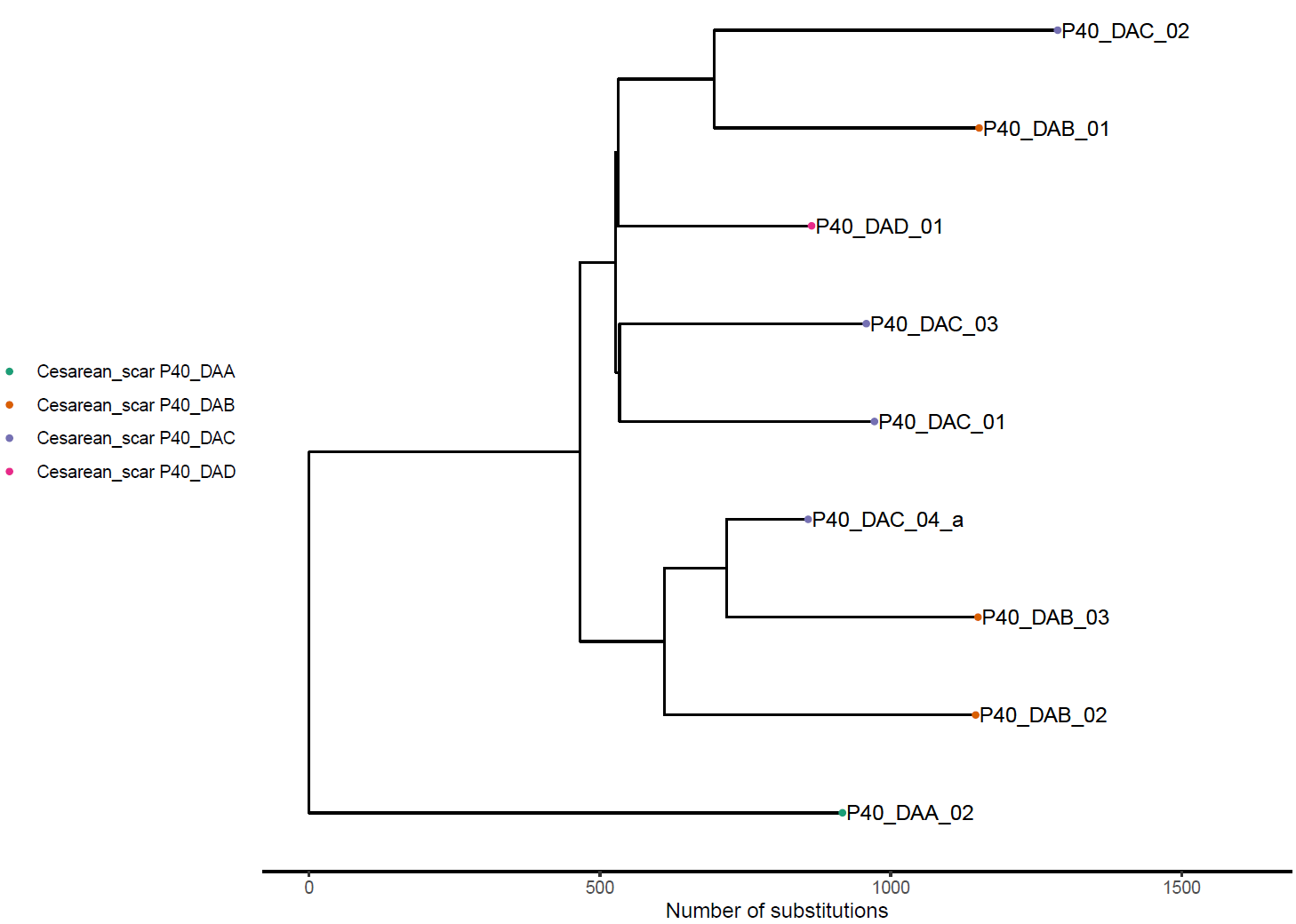

Fig. S25. Phylogenetic tree for Patient 41.

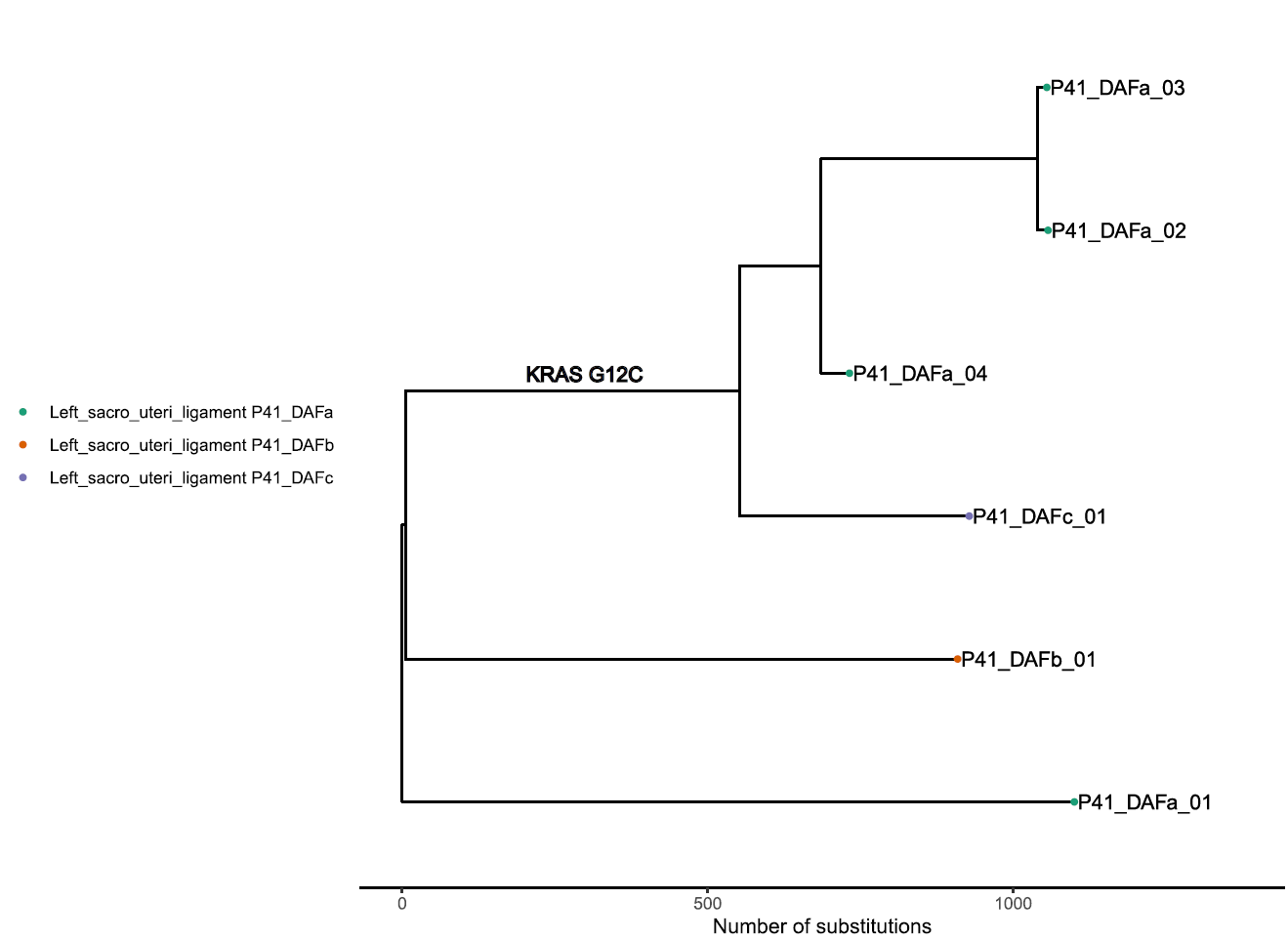

Fig. S26. Phylogenetic tree for Patient 42.

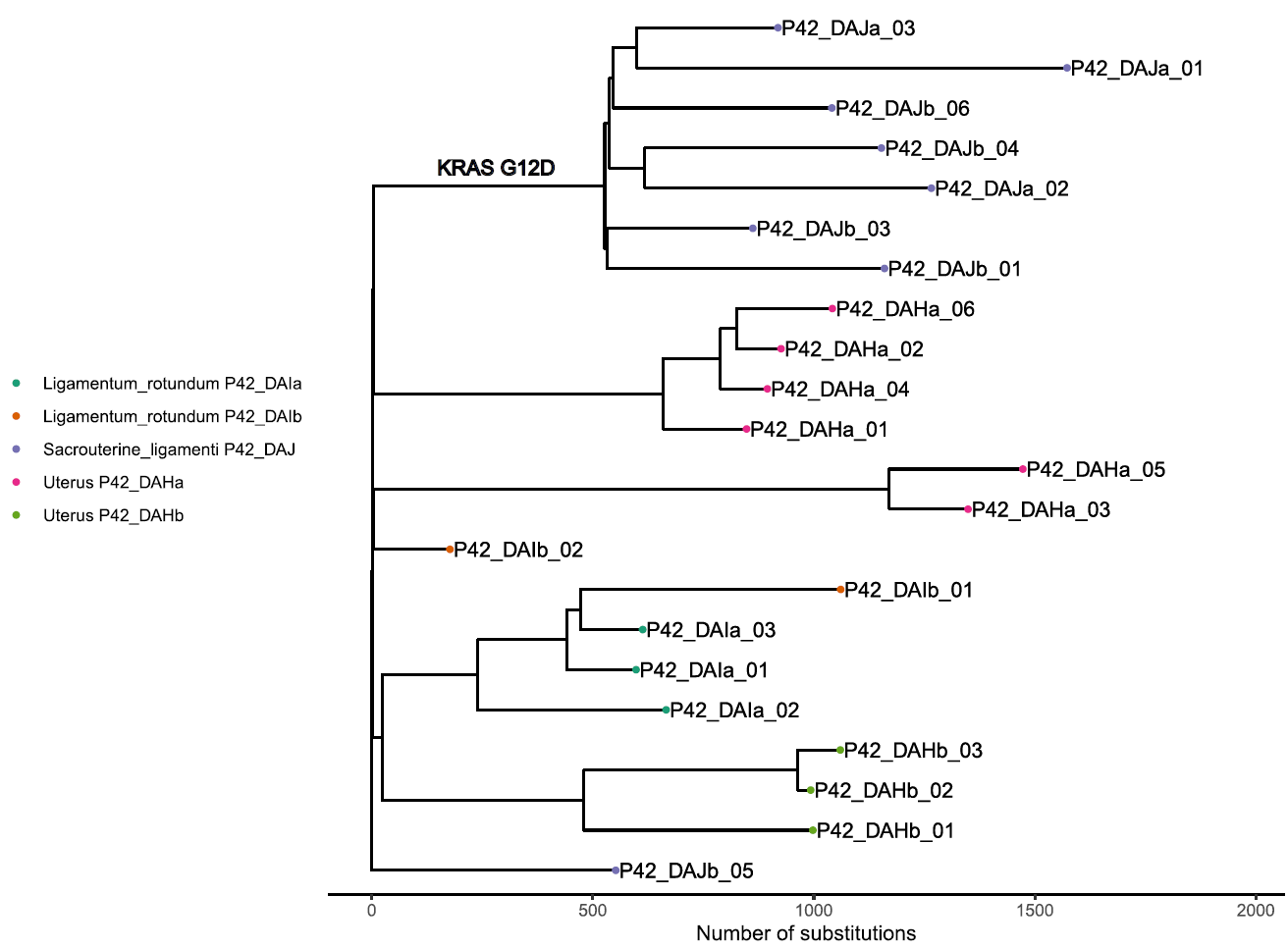

Fig. S27. Phylogenetic tree for Patient 43.

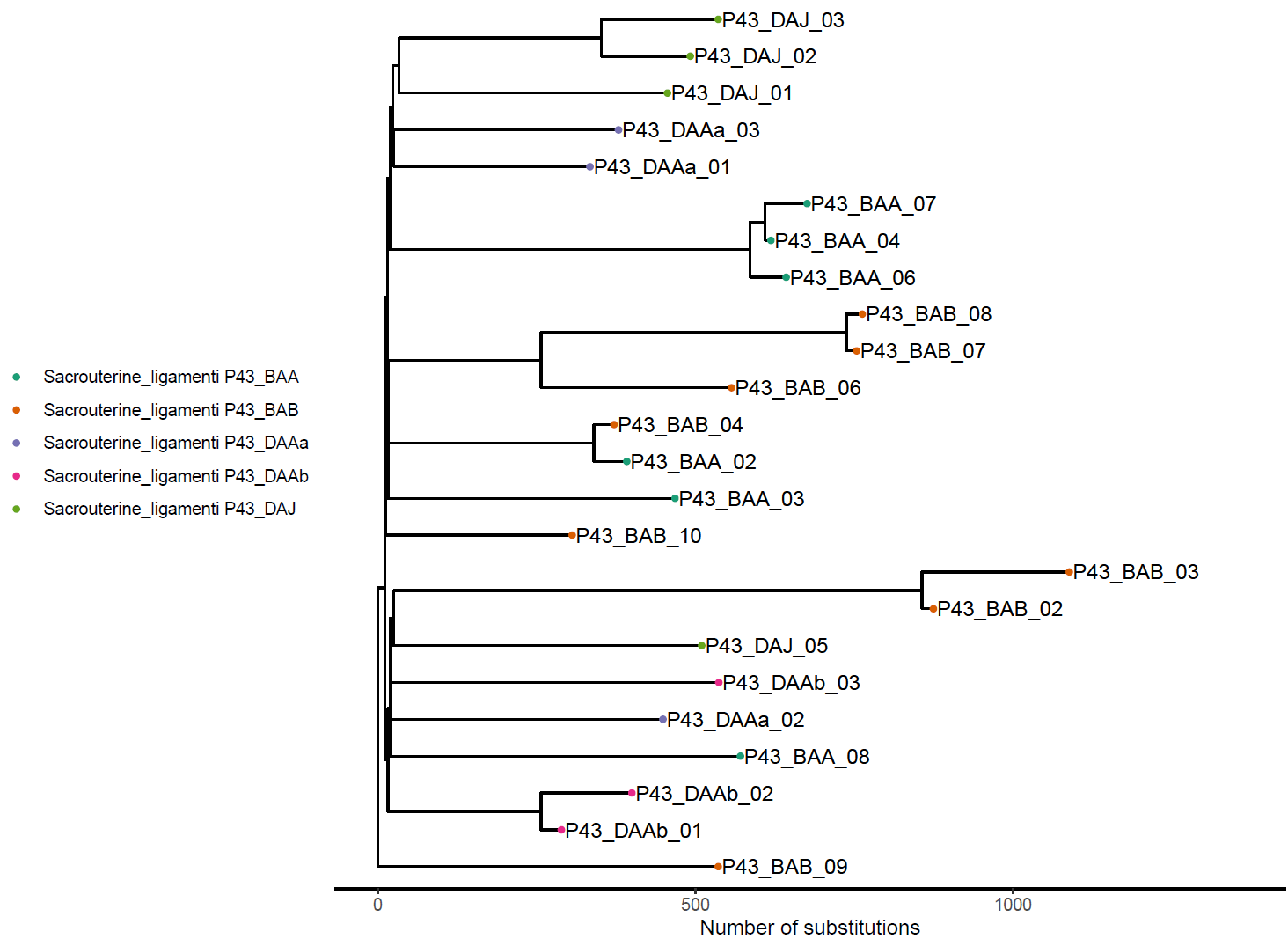

Fig. S28. Cell-of-origin analysis for endometriosis lesions stratified by lesion type.

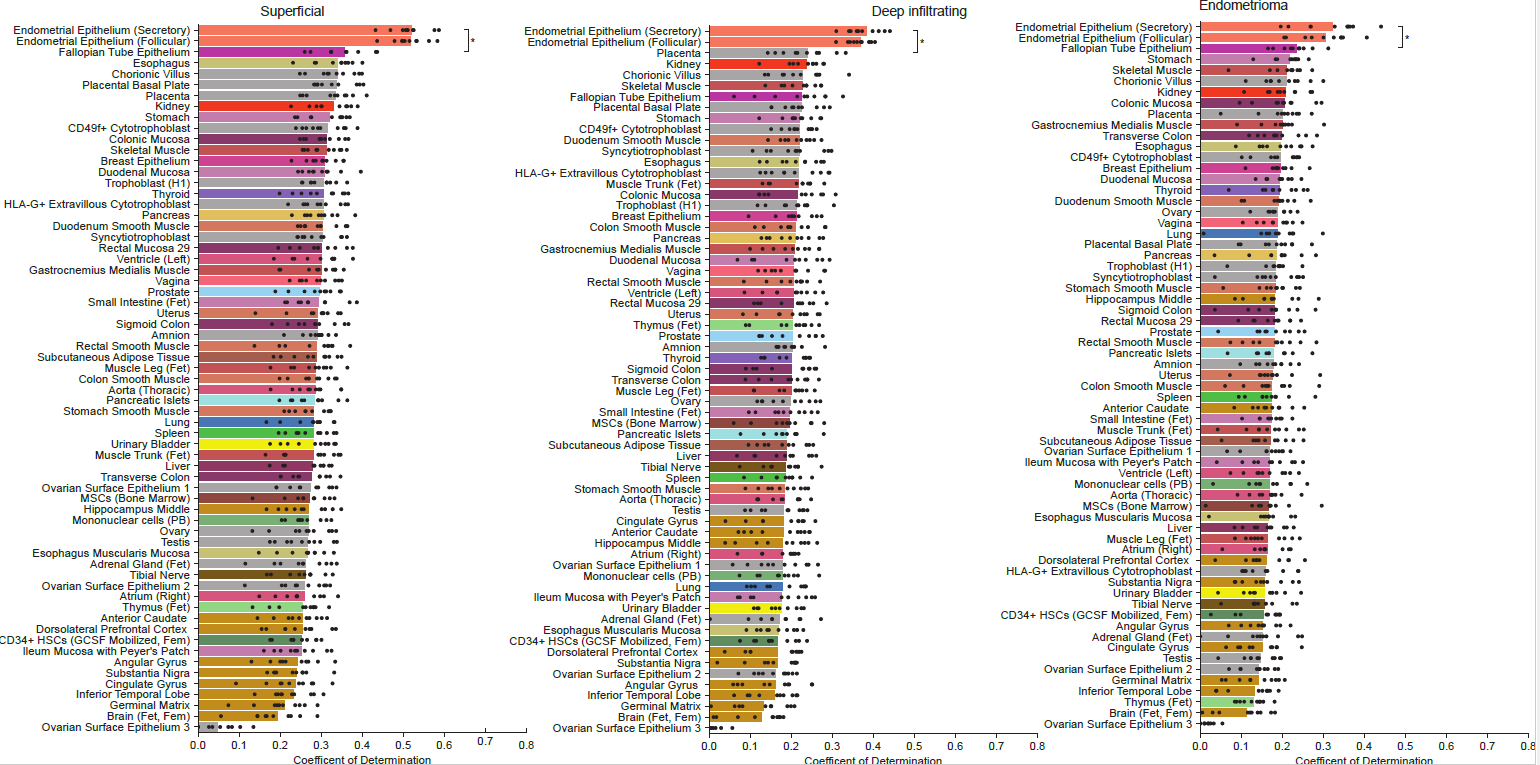

Performance of predicting mutational profiles from histone marks of 65 epigenomes representing distinct cell/tissue types; bars are colored according to tissue category; black points depict values obtained from 10-fold cross-validation. For every lesion type, normal endometrial cells are the best match.

Fig. S29. Cell-of-origin analysis for endometriosis lesions stratified by mutations that have occurred early vs late in molecular time.

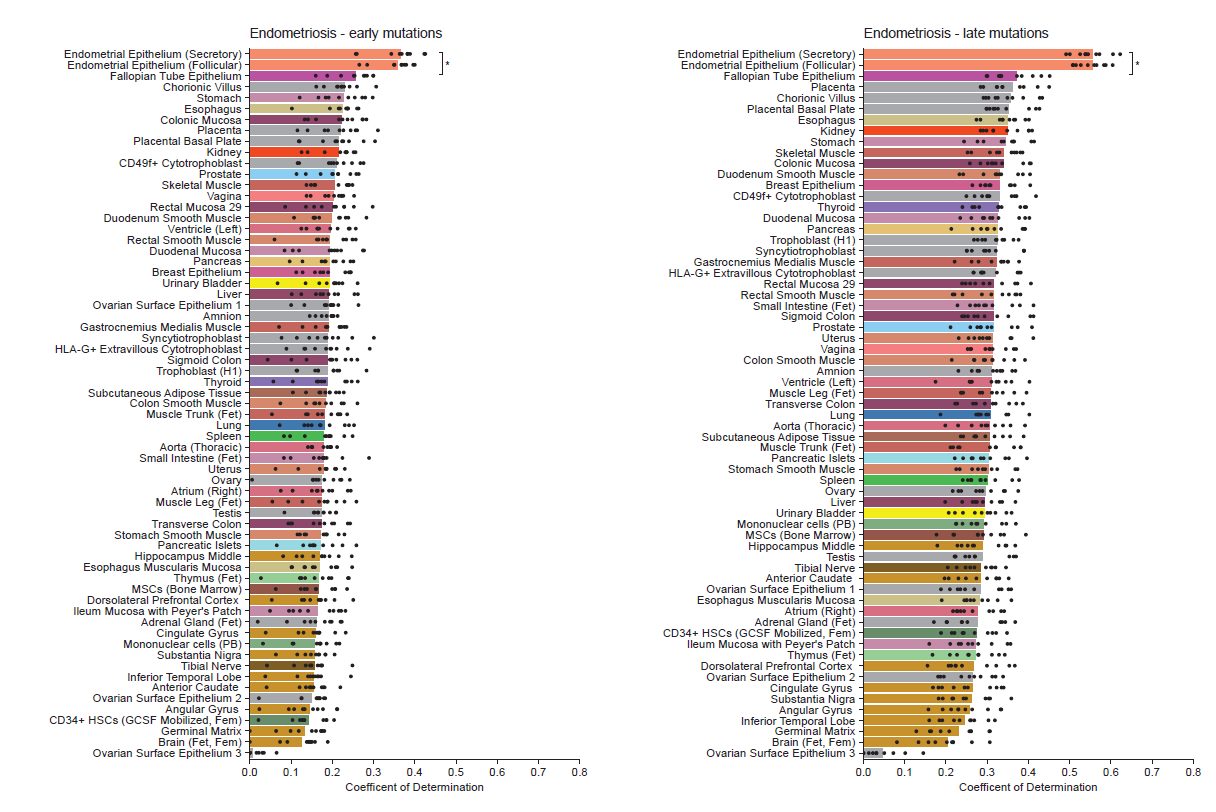

As described in section 9 of the method section above, mutations were defined as late-occurring if they followed a coalescent event that occurred after >20% of the molecular age of the clade-node as determined by the highest mutation burden of its children. The hypothesis was that if the metaplasia or circulating stem-cell theory were true, then the early mutations would be enriched in mutations reflecting the ancestral state. The bars represent the performance of predicting mutational profiles from histone marks of 65 epigenomes representing distinct cell/tissue types; bars are colored according to tissue category; black points depict values obtained from 10-fold cross-validation.

Captions for Supplementary Tables

**Table S1. Patient demographic information.**

TableS1_Patient_demographics.txt

**Table S2. Microbiopsy information.**

TableS2_Microbiopsy_metadata.txt

**Table S3. dNdScv selection analysis. A) Combined analysis across the whole cohort. B) Selection analysis restricted to mutations found in lesional samples. C) Selection analysis restricted to mutations found in normal samples.**

TableS3_dNdS_results.xlsx

**Table S4. Results of the Cell-of-origin analysis.**

TableS4_COO_results.txt

**Table S5. A list of mutations in *KRAS* and *PIK3CA* found in the screening of whole uteri.**

TableS5_KRAS_PIK3CA_mut_from_screen.xlsx

**Table S6. Primer sequences for PCR-based genotyping of endometrial segments at barcode sites.**

TableS6_Primer_sequences.xlsx
